# Circadian protein regulation in the green lineage I. A phospho-dawn of protein modification anticipates light onset in the picoeukaryote *O. tauri*

**DOI:** 10.1101/287862

**Authors:** Zeenat B. Noordally, Matthew M. Hindle, Sarah F. Martin, Daniel D. Seaton, T. Ian Simpson, Thierry Le Bihan, Andrew J. Millar

## Abstract

Diel regulation of protein levels and protein modification had been less studied than transcript rhythms. Here, we compare transcriptome data under light-dark cycles to partial proteome and phosphoproteome data, assayed using shotgun mass-spectrometry, from the alga *Ostreococcus tauri*, the smallest free-living eukaryote. 10% of quantified proteins but two-thirds of phosphoproteins were rhythmic. Mathematical modelling showed that light-stimulated protein synthesis can account for the observed clustering of protein peaks in the daytime. Prompted by night-peaking and apparently dark-stable proteins, we also tested cultures under prolonged darkness, where the proteome changed less than under the diel cycle. The dark-stable, prasinophyte-specific proteins were also reported to accumulate when *O. tauri* formed lipid droplets. In the phosphoproteome, 39% of rhythmic phospho-sites reached peak levels just before dawn. This anticipatory phosphorylation suggests that a clock-regulated phospho-dawn prepares green cells for daytime functions. Acid-directed and proline-directed protein phosphorylation sites were regulated in antiphase, implicating the clock-related, casein kinases 1 and 2 in phase-specific regulation, alternating with the CMGC protein kinase family. Understanding the dynamic phosphoprotein network should be facilitated by the minimal kinome and proteome of *O. tauri*. The data are available from ProteomeXchange, with identifiers PXD001734, PXD001735 and PXD002909. This submission updates a previous version, posted on bioRxiv on 4th April 2018, as https://www.biorxiv.org/content/10.1101/287862v1

**Highlight:** The phosphorylation of most protein sites was rhythmic under light-dark cycles, and suggested circadian control by particular kinases. Day-peaking, rhythmic proteins likely reflect light-stimulated protein synthesis in this microalga.

## Introduction

Responses to light are critical for organisms of the green lineage (Noordally and Millar, 2015; Paajanen *et al*., 2021). The rapid effects of photosynthetic light harvesting, for example on redox state and sugar metabolism, are complemented by signalling photoreceptors (Whitelam and Halliday, 2007) and the slower, 24-hour regulation by the biological clock (Millar, 2016; Creux and Harmer, 2019). Circadian regulation allows organisms to anticipate the predictable, day-night transitions of the diel cycle, complementing the responses to faster changes in light levels (Troein *et al*., 2011). Mehta et al. (2021) refer to these as ‘anticipatory’ and ‘reactive’ regulation. At the macromolecular level, the transcriptomes in the green lineage show widespread and overlapping regulation of mRNA abundance by both light and circadian signals (see below), whereas the diel regulation of proteins and their post-translational modifications had been less studied (Mehta *et al*., 2021). We addressed that gap using a minimal biological system, focussing on protein phosphorylation.

Phosphorylation of an existing protein is energetically inexpensive, occurs rapidly and can then alter protein activity through conformational change or intermolecular recognition (Khoury *et al*., 2011). These characteristics seem fitted to reactive regulation. Some plant photoreceptor proteins include protein kinases that initiate light signalling (Christie, 2007; Djouani-Tahri el *et al*., 2011*a*).

Protein synthesis is not only far slower but also among the costliest macromolecular processes (Scott *et al*., 2010; Karr *et al*., 2012), seemingly more suited to anticipatory regulation. Rhythmic regulation might then provide a selective advantage, loosely summarised as making proteins when they are needed in the diel cycle (Laloum and Robinson-Rechavi, 2022). That reasoning helped to interpret the co-regulation of functional clusters of RNAs, when transcriptome studies demonstrated that over 50% of Arabidopsis RNAs can be rhythmic under diel, light-dark cycles (LD) (Smith *et al*., 2004; Blasing *et al*., 2005; Michael *et al*., 2008). Most strikingly, almost the whole transcriptome of the marine unicellular alga *Ostreococcus tauri* was rhythmic in controlled conditions (Monnier *et al*., 2010) and this was also the most-rhythmic taxon among the diverse plankton of a Pacific timeseries (Kolody *et al*., 2019). The clock might also allow anticipation, to ensure that the proteins had been fully synthesised and assembled to their active state by the appropriate time.

Proteomic data, in contrast, revealed that most detected proteins had stable levels, with an average half-life >6 days in the model plant *Arabidopsis thaliana* (Li *et al*., 2017), suggesting little scope for diel rhythmicity. Timeseries under constant light or a diel cycle found up to 6% of rhythmic proteins (Baerenfaller *et al*., 2012, 2015)(Choudhary *et al*., 2016; Uhrig *et al*., 2021; Krahmer *et al*., 2022). The most short-lived, regulatory proteins are harder to detect, but such proteins seem to be exceptions to the general protein stability, consistent with mammalian systems (Doherty *et al*., 2009). Global regulation of protein synthesis is also clearly relevant in plants and algae (Piques *et al*., 2009; Juntawong and Bailey-Serres, 2012; Pal *et al*., 2013; Missra *et al*., 2015; Ishihara *et al*., 2015). In this context, circadian RNA regulation was proposed to offer a selective advantage through seasonal adaptation to day-length on a timescale of weeks (Seaton *et al*., 2018).

More protein phosphorylation sites change over the diel cycle, compared to protein levels (Kusakina and Dodd, 2012; Mehta *et al*., 2021). Protein phosphorylation in plants and algae is most directly light-regulated by the photoreceptor kinases (Christie, 2007; Djouani-Tahri el *et al*., 2011a), though light also affects the broader phosphoproteome (Turkina *et al*., 2006; Boex-Fontvieille *et al*., 2014; Schönberg *et al*., 2017), for example affecting 25% of Arabidopsis phosphopeptides within 30 minutes (Uhrig *et al*., 2021). Circadian studies in Arabidopsis under constant light found up to 23% rhythmic phosphopeptides (Choudhary *et al*., 2015; Krahmer *et al*., 2022). These studies suggest that light responses and the circadian clock in Arabidopsis each control five-to ten-fold more phosphopeptides than the diel rhythm of total protein level, so it is also important to understand which phospho-regulators mediate these effects.

The amino acid sequences of rhythmically-regulated phosphosites have implicated a range of protein kinases with overlapping contributions in Arabidopsis (Choudhary *et al*., 2015; Uhrig *et al*., 2021; Krahmer *et al*., 2022). However, ∼1000 protein kinases shape the phosphoproteome in Arabidopsis (Champion *et al*., 2004) including several in plastids (Baginsky and Gruissem, 2009), compared to half that number in the human genome (Manning *et al*., 2002). Of particular interest, the casein kinases (CK1, CK2) and Glycogen Synthase Kinase 3 (GSK3), affect the circadian timing of all organisms suitably studied (Mehra *et al*., 2009). These kinases have central positions in the yeast kinase-target network (Breitkreutz *et al*., 2010) and are highly conserved (Hindle *et al*., 2014), in contrast to photoreceptor proteins or circadian transcription factors (Noordally and Millar, 2015; Dunlap and Loros, 2017).

Here, we compare the prevalence of proteomic and phosphoproteomic regulation under LD cycles, using *O. tauri* as a minimal model for the green lineage (Noordally and Millar, 2015). This alga not only has a ubiquitously-rhythmic transcriptome, but its genome is also reduced to 13Mbp (Blanc-Mathieu *et al*., 2014), likely due to selection pressure to reduce cell size to 1-2µm (Courties *et al*., 1994). Its 7699 protein-coding genes include just 133 protein kinases that represent the core families for eukaryotic signalling (Hindle *et al*., 2014) and a minimal set of Arabidopsis clock gene homologues (Corellou *et al*., 2009; Djouani-Tahri el *et al*., 2011*b*; Troein *et al*., 2011; Ocone *et al*., 2013). CK1 and CK2 modulate circadian timing in the light, with widespread effects on the algal phosphoproteome (Le Bihan *et al*., 2011, 2015; van Ooijen *et al*., 2013). A non-transcriptional, 24-hour oscillator of unknown mechanism was also revealed under prolonged darkness, when transcription stops in this organism (O’Neill *et al*., 2011; van Ooijen *et al*., 2011; Edgar *et al*., 2012; Bouget *et al*., 2014; Feeney *et al*., 2016). In cyanobacteria, the non-transcriptional clock is driven by rhythmic protein phosphorylation, so rhythmic protein kinase activities could also be relevant in *O. tauri* (van Ooijen and Millar, 2012; Wong and O’Neill, 2018).

Our results reveal widespread daily rhythms in both the proteome and phosphoproteome in *O. tauri*, including expected features such as the diel control of conserved, cell cycle phospho-regulators. Rather than rapid phosphorylation responses and slow, rhythmic anticipation in protein profiles, however, the level of many rhythmic proteins appears light-responsive, whereas much of the rhythmic phosphoproteome anticipates dawn. The phosphosite sequences strongly implicate phase-specific protein kinase classes. Moreover, we identify a set of rhythmic, algal-specific proteins that accumulate in prolonged darkness and were also identified in conditions that promote the formation of lipid droplets.

## Materials and Methods

### Materials

Chemicals were purchased from Sigma-Aldrich (now a subsidiary of Merck Life Science UK Ltd, Dorset, UK) unless otherwise stated. Main solvent, acetonitrile and water for liquid chromatography– dual mass spectrometry (LC-MSMS) and sample preparation were HPLC quality (Thermo Fisher Scientific, Loughborough, UK). Formic acid was Suprapure 98-100% (Merck) and trifluoroacetic acid (TFA) was 99% purity sequencing grade. Porcine trypsin TPCK treated was from Worthington (Lorne Laboratories, Reading, UK). All HPLC-MS connectors and fittings were from Upchurch Scientific (Hichrom, Theale, UK) or Valco (RESTEK, High Wycombe, UK). % are expressed in v/v.

### *O. tauri* media and culturing

*Ostreococcus tauri* OTTH95 were cultured as previously described (van Ooijen *et al*., 2012), supplemented with 0.22 μm filtered 50 µg ml^-1^ ampicillin, neomycin and kanamycin antibiotics in vented tissue culture flasks (Sarstedt, Leicester, UK). Cultures were maintained by splitting weekly at 1:50 dilution. In preparation for proteomics experiments, cultures were grown in growth media supplemented with 200 mM sorbitol and 0.4% glycerol prior to harvesting (O’Neill *et al*., 2011). Cells were cultured under cycles of 12 hour light/ 12 hour dark (LD) at 20°C in a controlled environment chamber (MLR-350, Sanyo Gallenkamp, Loughborough, UK) at a light intensity of 17.5 μEm^-2^ s^−1^ white fluorescent light filtered by 724 Ocean Blue filter (LEE Filters Worldwide, Andover, UK).

### *O. tauri* cell harvesting

Cells were grown for 7 days in LD and on the seventh day harvested, with five replications, at Zeitgeber Time (ZT) 0, 4, 8, 12, 16 and 20, where ZT0 corresponds to dawn. At ZT0 cells were harvested a few minutes before the lights went on and at ZT12, before the lights went off. 135 ml culture was harvested by centrifugation (4000 rpm, 10 min, 4°C) per sample replicate, each from a separate culture vessel. Pellets were resuspended in ice cold phosphate buffered saline solution (PBS). Cultures were centrifuged as before, pellets were air dried and then vortex-mixed in 250 µl 8M urea and stored at -80°C. For total cell lysate, cells were dissolved by sonication (Branson Ultrasonics) and diluted with 500 µl dH_2_O. Cells were grown for 7 days in LD and on the eighth day the Dark Adaptation (DA) experiment cell harvests were performed at ZT24, 48, 72 and 96 in constant darkness with five replications. The samples were harvested and prepared as for the LD experiment.

### Protein digestion

Samples were analysed by Bradford Assay (Bio-Rad, Watford, UK) and 400 µg protein of each sample was used in the digestion. Samples were reduced in 10 mM dithiothreitol and 50 mM ammonium bicarbonate, and alkylated with 25 mM iodoacetamide. Samples were digested overnight with 10 µg (1:40 ratio) trypsin under agitation at room temperature at pH8 in a total volume of 1 ml. Samples were cleaned on SPE BondElut 25 mg columns (Agilent Technologies, Stockport, UK) following the vendor instruction. 50 µl (∼20 µg) was removed and dried for LC-MS (Speedvac, Thermo Fisher Scientific). The remaining ∼380 µg were also dried in preparation for phosphopeptide enrichment, and stored at -20°C.

### Phosphopeptide enrichment

Dried peptide samples (∼380 µg) were sonicated in 50 µl solution 0 (2.5% acetonitrile, 0.5% TFA) and 100 µl solution 2 (80% acetonitrile, 0.5% TFA, 100% lactic acid). Titansphere Phos-TiO Kit spin tip-columns (GL Sciences, Tokyo, Japan) were washed with 40 µl solution 1 (80% acetonitrile, 0.5% TFA). Samples were loaded on the spin tip-columns and passaged three times through a centrifuge; 5 min at 200 xg, 15 min incubation at room temperature and 10 min at 200 xg. Spin tip-columns were subsequently washed once with solution 1, twice with solution 2 and twice with solution 1for 2 min at 200x g. Phosphopeptides were eluted in two steps, first with 50 µl 5% ammonium hydroxide (5 min at 200 xg) and secondly, with 5% pyrrolidine solution. 20 µl 20% formic acid was added to lower the pH and samples were cleaned on Bond Elut OMIX C18 pipette tips (Agilent Technologies) following the manufacturer’s instruction.

### Protein and phosphoprotein quantification

15 µg protein from total *O. tauri* cell lysates were run on a Novex NuPAGE 4-12% Bis-Tris by SDS-PAGE with PeppermintStick Phosphoprotein Molecular Weight Standards and Spectra Multicolor Broad Range Protein Ladder (Thermo Fisher Scientific). The gel was fixed overnight (50% methanol, 40% ddH_2_O, 10% glacial acetic acid), washed in ddH_2_O and stained with Pro-Q Diamond Phosphoprotein Gel Stain (Invitrogen, now Theremo Fisher Scientific, Loughborough, UK) in the dark at 25°C following manufacturer’s instructions. The gel was imaged on a Typhoon TRIO variable mode imager (GE Healthcare, Amersham, UK) at 532 nm excitation/ 580 nm emission, 450 PMT and 50 micron resolution. Images were processed using ImageQuant TL software (GE Healthcare, Amersham, UK). The gel was re-used for protein quantification using SYPRO Ruby Protein Gel Stain (Themo Fisher Scientific, Loughborough, UK) following manufacturer’s instructions and imaged using a UV transilluminator (Ultra-Violet Products Ltd, Cambridge UK). Protein and phosphoprotein bands were quantified using Image Studio Lite v 4.0 (LI-COR Biosciences, Cambridge, UK).

### Protein per cell quantification

Cells were grown (as described above) and independent, triplicate cultures were harvested at the times indicated. Cultures were monitored using spectrophotometry at 600nm. Total protein was quantified using the Quick Start Bradford Assay following manufacturer instructions (Bio-Rad, Watford, UK). Cell number was estimated either by counting four fields of view per culture in a haemocytometer after trypan blue staining (Abcam protocols, Cambridge, UK), or by fluorescence-activated cell sorting (FACS). For FACS, a 1/200 dilution of cells were transferred to fresh media containing 1X SYBR Green I Nucleic Acid Gel Stain (Invitrogen, now Theremo Fisher Scientific, Loughborough,UK) and FACS-counted (FACScan, BD Bioscience, Wokingham, UK) at a flow rate of 60μl per minute.

### qPCR for transcriptional regulation during dark adaptation (DA)

Cells were cultured and harvested in the same experimental regime (described above) and harvested in biological triplicate at the times indicated for the LD and DA experiments. Total RNA was extracted from frozen cells using an RNeasy Plant Mini Kit and DNase treated (QIAGEN, Manchester, UK). First-strand cDNA was synthesised using 1 µg RNA and 500 ng µl^-1^ Oligo(dT)_15_ primer (Promega, Southampton, UK), denatured at 65°C for 5 min, and reverse transcribed using SuperScript II (Invitrogen, now Theremo Fisher Scientific, Loughborough, UK) at 42 °C for 50 min and 70 °C for 10 min. 1/100 cDNA dilutions were analysed using a LightCycler^®^480 and LightCycler^®^480 SYBR Green I Master (Roche, Welwyn Garden City, UK) following manufacturer’s instructions and cycling conditions of pre-incubation 95°C for 5 min; 45x amplification cycles of 95°C for 10 s, 60°C for 10 s, 72°C for 10 s. The following 5’ to 3’ forward (F) and reverse (R) primers to *O. tauri* gene loci were used: ostta01g01560 GTTGCCATCAACGGTTTCGG (F), GATTGGTTCACGCACACGAC (R); ostta03g00220 AAGGCTGGTTTGGCACAGAT (F), GCGCTTGCTCGACGTTAAC (R); ostta03g04500 GCCGCGGAAGATTCTTTCAAG (F), TCATCCGCCGTGATGTTGTG (R); ostta04g02740 ATCACCTGAACGATCGTGCG (F), CCGACTTACCCTCCTTAAGCG (R); ostta10g02780 GGCGTTCTTGGAATCTCTCGT (F), TATCGTCGATGATCCCGCCC (R); ostta10g03200 GGTACGGAGGAAGAAGTGGC (F), ATGTCCATGAGCTTCGGCAA (R); ostta14g00065 GACAGCCGGTGGATCAGAAG (F), TCGAGGTAGCTCGGGAGATC (R); ostta16g01620 ACGGGTTGCAGCTCATCTAC (F), CCGCTTGGGTCCAGTACTTC (R); ostta18g01250 CTTGCAAATGTCCACGACGG (F), ATGATGTGGCACGTCTCACC (R); OtCpg00010 ACATGACTCACGCGCCTTTA (F), TGCCAAAGGTGCCCTACAAA (R). Primers to eukaryotic translation elongation/initiation factor (EF1a) ostta04g05410 GACGCGACGGTGGATCAA (F) and CGACTGCCATCGTTTTACC (R) were used as an endogenous control. Data were combined for biological and two technical replicates and relative quantification performed using LightCycler^®^480 1.5 software (Roche).

### HPLC–MS analysis

Micro-HPLC-MS/MS analyses were performed using an on-line system consisting of a micro-pump 1200 binary HPLC system (Agilent Technologies) coupled to an hybrid LTQ-Orbitrap XL instrument (Thermo Fisher Scientific). The complete method has been described previously (Le Bihan *et al*., 2010). For all measurements, 8µl of sample was injected using a micro-WPS auto sampler (Agilent Technologies) at 5µl /min. After sample loading, the flow rate across the column was reduced to approximately 100-200 nl/min using a vented column arrangement. Samples were analysed on a 140 min gradient for data dependant analysis.

### HPLC-MS data analysis

To generate files compatible with public access databases PRIDE (Vizcaino *et al*., 2016) and the former pep2pro (Hirsch-Hoffmann *et al*., 2012), Mascot Generic Format (MGF) input files were generated using MSConvert from ProteoWizard (Kessner *et al*., 2008). MSMS data was searched using MASCOT version 2.4 (Matrix Science Ltd, London, UK) against the *O. tauri* subset of the NCBI protein database (10114 sequences from NCBI version 2014 June 6th including common contaminants) using a maximum missed-cut value of 2, variable oxidation (M), N-terminal protein acetylation, phosphorylation (STY) and fixed carbamidomethylation (C); precursor mass tolerance was 7 ppm and MSMS tolerance 0.4 amu. The significance threshold (p) was set below 0.05 (MudPIT scoring). Global FDR was evaluated using decoy database search and removal of peptides ranked higher than 1 for a mascot score above 20 (∼1% global FDR). Mass spectrometry proteomics data have been deposited in PRIDE ProteomeXchange Consortium (Vizcaino *et al*., 2014) via the PRIDE partner repository with the dataset identifier LD global proteomics, PXD001735; LD phosphoproteomics, PXD001734; DA global proteomics, PXD002909. Data was converted into PRIDEXML using Pride converter 2.0.20 and submitted using proteome exchange tool pxsubmission tool 2.0.1. The LC-MS data were also publicly available in the former pep2pro database (Assemblies ‘Ostreococcus tauri Light:dark cycle,LD global’, ‘Ostreococcus tauri Light:dark cycle,LD phospho’, and ‘Ostreococcus tauri dark adaptation,DA global’). Label-free quantification was performed using Progenesis version 4.1 (Nonlinear Dynamics, Newcastle, UK). Only MS peaks with a charge of 2+, 3+ or 4+ and the five most intense spectra within each feature were included in the analysis. Peptide abundances were mean-normalised and ArcSinH transformed to generate normal datasets. Within-group means were calculated to determine fold changes. Neutral losses of phosphoric acid typical of serine and threonine phosphorylated were validated manually in all significantly differential phosphopeptides. Ambiguous sites were confirmed by cross-referencing (by sequence, charge, and quantity of residue modifications) with most probable site predictions from MaxQuant version 1.0.13.8 (Cox and Mann, 2008) in singlet mode, Mascot settings as above. Where multiple occurrences of residue phosphorylation events were quantified, abundances were summed, collating all charge states, missed cuts and further modifications.

### Data analysis

#### Merging

For accurate and unique phosphopeptide quantification we addressed variant redundancy at different charge states, alternative modifications (e.g. oxidation and acetylation) and multiple sites of protease digestion. All unique phosphorylation events were retained, including multiple phosphorylation, at a given amino acid motif, while summing the quantification of these technical variants. The qpMerge (http://sourceforge.net/projects/ppmerge/) software was used to combine Progenesis and MaxQuant phospho-site predictions and produce a unique set of quantified phosphopeptide motifs (Hindle *et al*., 2016).

#### Outlier identification and removal

To detect outliers we applied principal component analysis (PCA) of replicates and comparing each replicates *r*^*2*^ to the respective median abundance at that ZT. A single replicate, 4E, was excluded based on extreme differences in peptide quantification as viewed in the wider distribution of the ratios to the sample median, which was confirmed with a Pearson’s correlation against a defined criterion of a sample median of < 0.8 (Supplementary Figures S1).

#### P-value calculation and false discovery rate (FDR)

For analysing the significance of changing protein and peptide abundance over time, non-linear response of expression using polynomial regression was modelled using the R Stats Package. A third order polynomial was fitted, allowing for an expected peak and trough within a 24 h daily cycle. An arcsinh transformation of abundance was applied to meet the required assumption of normality (Burbidge *et al*., 1988). FDR was calculated using the Benjamini and Hochberg (BH) method (Benjamini and Hochberg, 1995). More than 2 quantifying peptides were required to report protein abundance.

#### Equivalence testing

Using the R equivalence package, the statistical equivalence of mean abundance across time was tested as the highest *p*-value from exhaustive pairwise Two one-sided test approach (TOST) tests over all ZTs (Schuirmann, 1981; Westlake, 1981). We tested whether abundances had upper and lower differences of less than 0.3 within the equivalence margin (ε).

#### O. tauri gene identifiers

*O. tauri* genome version 1 gene IDs (Derelle *et al*., 2006) for microarray data were converted to version 2 IDs (Blanc-Mathieu *et al*., 2014) by finding exact sequence matches for the microarray probes (Accession GPL8644) (Monnier *et al*., 2010) in the version 2 FASTA coding sequence file.

#### Principle component analysis (PCA)

PCA was used to investigate the main components of variation in the data using prcomp from the R Stats Package. The abundances were zero-centred per-feature. The PCA values for each feature were extracted and then used for Gene Ontology (GO) enrichment analysis.

#### Clustering

Hierarchical clustering was performed with hclust from the R Stats Package and applied on per-feature (protein or phosphopeptide motif) mean abundances over time, which were zero-centred and scaled. Pearson’s correlation was used to calculate distance matrix and the Ward method (Ward, 1963) for linkage criteria. The hierarchical tree was divided into clusters using the dynamicTreeCut algorithm (Langfelder *et al*., 2008). The hybrid cut tree method with a cut height of 100 and a minimum cluster size of 20 was used for both datasets. Clusters are displayed with 95% (black lines) and 99% (orange lines) confidence via multiscale bootstrap resampling (AU determined p-value).

#### Enrichment analysis for GO terms

TopGO was used to evaluate the enrichment of GO terms, for each ontology aspect, within clusters, peaks, troughs, and principal components. For clusters, peaks and troughs a Fisher’s exact test was used by partitioning at 95% confidence on FDR corrected *p*-values, and with a fold change >1.5 in normalised abundance. For each test, we use a relevant background of non-significant observed features. To test for enrichment of GO terms for each PCA the Kolmogorov-Smirnov test was applied over the absolute PCA values for each gene. GO terms were predicted by InterProScan 5 (Jones *et al*., 2014) on amino acids sequences for *O. tauri* coding sequences (NCBI version 140606 (Blanc-Mathieu *et al*., 2014)).

#### Homology modelling

Structural homology models were generated using I-TASSER (Yang and Zhang, 2015) for prasinophyte-family specific proteins of unknown structure and function, including for ostta02g03680 compared to the human Bar-domain protein structure in PDB entry with DOI 10.2210/pdb2d4c/pdb. Other suggested homologies were more limited.

#### pLOGO and binomial statistics

Significantly over- and under-represented amino acid residues at different time-points were calculated using the binomial based pLogo tool (O’Shea *et al*., 2013). The Motif-X tool (Chou and Schwartz, 2011) was used to discover novel motifs in the dataset. Binomial statistics were applied to calculate the enrichment of motifs and the combined probabilities of amino acids with similar properties in a phospho-motif (*e*.*g*. the acidic D/E positions in the CK2 motif).

#### Kinase target prediction

Computational prediction of protein kinase motifs associated with the identified phosphorylation sites was performed using Group-based Prediction System, GPS Version 3.0 (http://gps.biocuckoo.org/index.php) (Xue *et al*., 2011).

#### *O. tauri* loci IDs mapping to *A. thaliana* loci IDs

*O. tauri* and *A. thaliana* IDs were mapped using EggNOG4.1 (http://eggnogdb.embl.de). *O. tauri* proteins were downloaded from https://bioinformatics.psb.ugent.be/gdb/ostreococcusV2/LATEST/OsttaV2_PROT_20140522.fasta.gz (May 22^nd^, 2014). Viridiplantae (virNOG) hmms and their descriptions and annotations were transferred to *O. tauri* proteins using hmmr 3.1 (http://hmmer.janelia.org)

### Mathematical simulations

#### Simulated protein rhythms

Protein dynamics (*P(t)*) were simulated according to the following model:

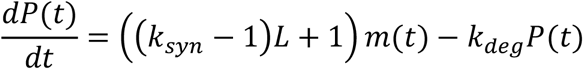

Where *L(t)* = 1 during the day (ZT <=12), and 0 otherwise. The rate of protein degradation (*k*_*deg*_) was set to 0.1 h^-1^, and the ratio of protein synthesis in the light compared to the dark (*k*_*syn*_) was set to 4, based on (Martin *et al*., 2012). The rhythmically expressed mRNA levels (*m(t)*) are given by:

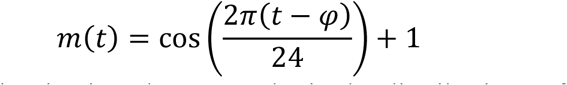

The peak phase of expression is given by *φ*. To obtain the distributions of peak and trough protein levels, the peak phases (*φ*) of mRNA expression were uniformly distributed at 0.1 h intervals across the range [0,24]. For each phase of mRNA expression, the timing of peak and trough protein levels was determined by simulating the model dynamics in MATLAB using the ode15s ODE solver. The peaks and troughs were identified across a 24 h period, following 240 h simulation to allow the dynamics to reach a steady behaviour (i.e. with the same protein levels at ZT0 and ZT24).

#### Protein degradation rates and depletion during dark adaptation

Degradation rates were calculated from published proteomics data (Martin *et al*., 2012), which characterised the dynamics of partial ^15^N isotope incorporation. We assumed a labelling efficiency of 0.93 (=maximum labelled fraction achieved of any protein + 0.01), and fitted a simple kinetic model assuming: (1) constant labelling efficiency over time; (2) different proteins are labelled at the same efficiency; (3) heavy and light fractions are turned over at equal rates, similar to (Seaton *et al*., 2018). One protein with a high degradation rate ∼0.03 h^-1^ was excluded as an outlier, which increased the correlation from R = -0.48 to -0.7 when included.

## Results

To understand the landscape of protein abundance and phosphorylation across the diel cycle, we harvested quintuplicate biological samples of *O. tauri* at six timepoints across a 12 h light/12 h dark (LD) cycle. Dawn samples (zeitgeber time 0, ZT0) were harvested just before lights-on, and samples at ZT12 before lights-off, to detect biological regulation that anticipated these transitions. The proteome and phosphoproteome were measured in whole-cell extracts from each sample, by label-free, liquid chromatography–mass spectrometry (Figure 1A). After removing a technical outlier (Supplementary Fig. S1), 855 proteins were quantified with 2 or more peptides (Supplementary Table S1). Phosphopeptides were enriched by metal-affinity chromatography prior to detection. For quantification, we combined the phosphopeptide species that shared phosphorylation on a particular amino acid, irrespective of other modifications (Hindle *et al*., 2016). We refer to this set of phosphorylated species as a phosphopeptide motif (PM). 1472 phosphopeptide motifs were quantified, from 860 proteins (Supplementary Table 2). Serine and threonine residues were modified most; only 1% of PMs included phospho-tyrosine. The quantified proteins and phosphoproteins each represent ∼11% of the total *O. tauri* proteome (Figure 1B). 29 out of 61 proteins encoded on the chloroplast genome (Robbens *et al*., 2007) were quantified, with 6 PMs. 3 out of 43 mitochondrial-encoded proteins were quantified with no PMs, consistent with other studies (Ito *et al*., 2009).

**Figure 1.**
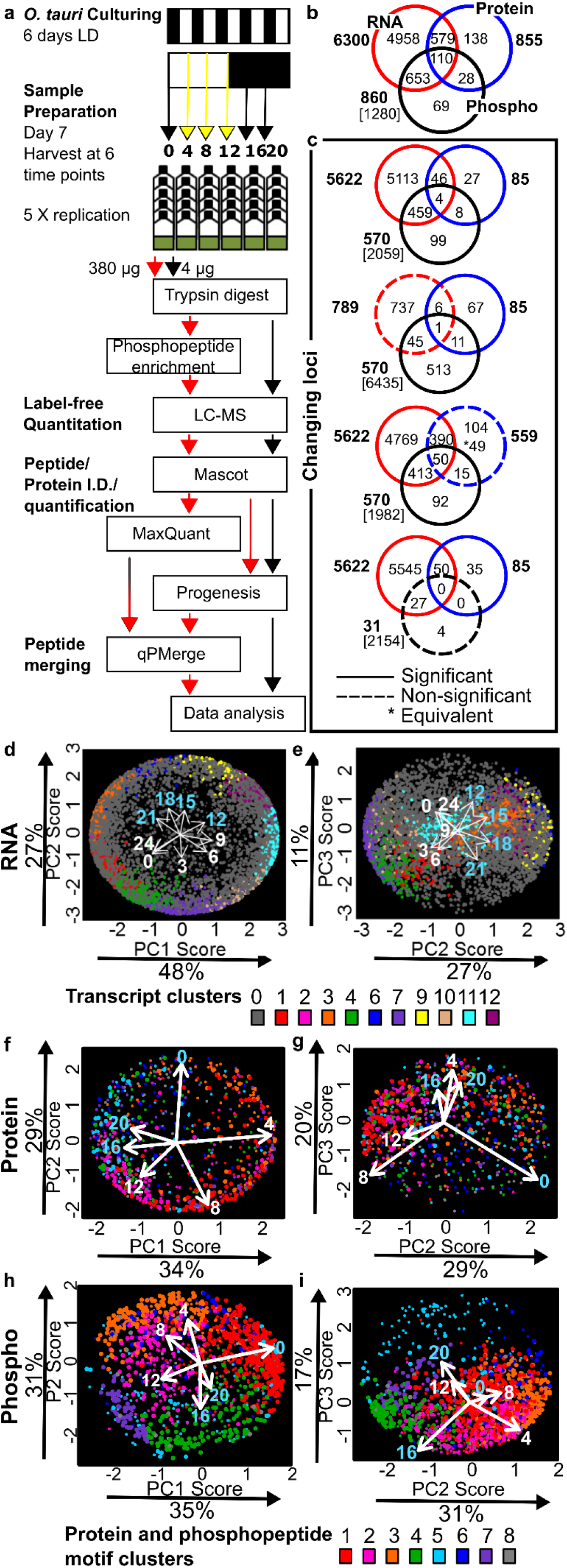
Daily variation in transcripts, proteins, and phosphopeptide motifs. (A) Workflow for proteomics in *O. tauri* under LD. Overlap in (B) detected and quantified gene loci, (C) significantly changing (solid circles) or not significantly-changing (dashed circles) loci for transcripts (Monnier *et al*., 2010), proteins and PMs; genomic loci excluded (square brackets). (D-I) Bi-plots of PCA for (D, E) transcript, (F, G) protein and (H, I) phosphomotif profiles. Proportion of the variance for each PC is indicated. Dot locations show the weighting of each RNA/protein/PM in each PC; colours show the assigned cluster (as in Supplementary Figure S4).

### Diel rhythmicity of the transcriptome, proteome and phosphoproteome

To compare the patterns and prevalence of daily rhythms at different regulatory levels, we re-analysed published transcriptome data in parallel with these protein and phosphoprotein data, summarised in Figure 1C. Gene expression in *O. tauri* was strongly rhythmic under LD cycles, with 89% of transcripts scored rhythmic, as previously reported (Monnier *et al*., 2010). 85 (9.5%) of the detected proteins were significantly rhythmic and changed by at least 1.5-fold, with only 11 of these proteins changing level by more than 5-fold. In contrast, 66% of phosphoproteins or 58% of PMs (570 of 860 proteins; 850 of 1472 PMs) were rhythmic by these criteria and the levels of 35 PMs changed more than 20-fold. The overlap among all three datasets included only 110 genes. The most common pairwise overlaps involved genes with changing levels of RNA and/or PMs but not of protein (Figure 1C).

Protein levels nonetheless changed smoothly, with distinct waveforms. Of the twenty most highly-detected proteins, likely including the most abundant, 11 were significantly rhythmic but with low amplitudes (Supplementary Figure S2A), such that only ostta10g03200 exceeded the 1.5-fold change threshold (Table S1). 15 of the twenty most highly-detected PMs, in contrast, were rhythmic by both criteria (Supplementary Figure S2B). The more stringent, “equivalence” test revealed 49 proteins with significantly non-changing protein abundance but with significantly changing transcript and PMs, illustrated by the 10-fold change in PM abundance on the non-changing, chlorophyll-binding protein CP26, amongst others (Supplementary Figure S3).

To address our major question on the dominant patterns of regulation, we used undirected, principal component (PC) analysis (Fig. 1D-1I). Clustering (Fig. 1D-1I, Supplementary Figure S4) and analysis of peak distributions (Fig. 2A-C) informed more detailed hypotheses on upstream regulation and downstream, functional effects. The PC analysis represented most (83-86%) of the variance in the data sets but indicated a differing balance of molecular regulation (Supplementary Table S3). The transcriptome and phosphoproteome data clearly separated between dawn and dusk timepoints (in PC1), whereas the light and dark intervals were separated by the secondary PC2. This mapped the 13 transcriptome and 6 phosphoproteome timepoints into their respective, temporal sequences. Gene Ontology (GO) terms relating to translation, ribosome biogenesis and RNA processing were enriched among dawn-expressed RNAs, and mitotic processes (DNA replication and repair) among dusk-expressed transcripts (Supplementary Table S3), similar to past analysis (Monnier *et al*., 2010). The functions of rhythmic phosphoproteins are discussed in more detail, below.

**Figure 2.**
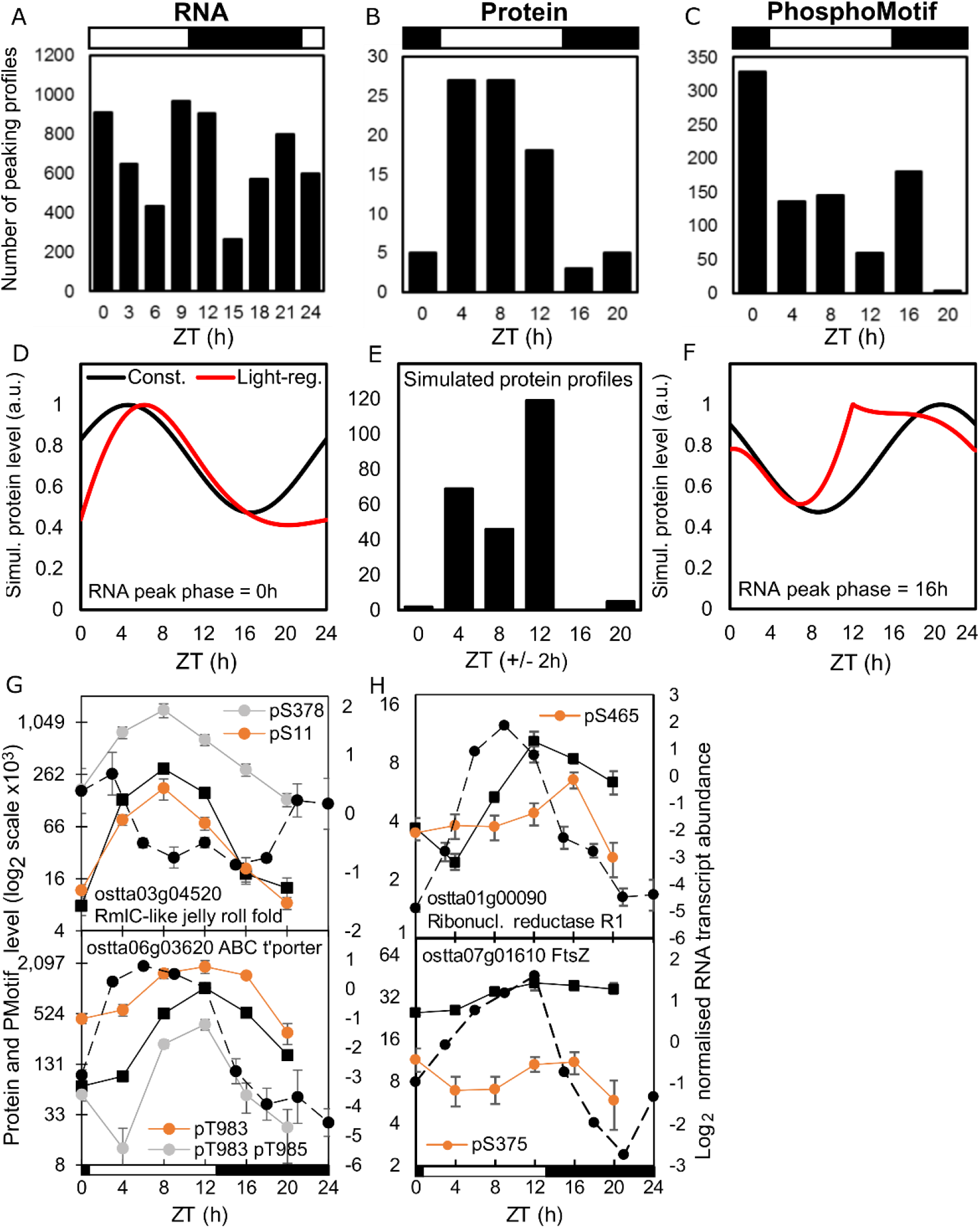
Distribution of rhythmic protein and phosphopeptide motif peaks, with examples. Temporal distribution of peaking profiles in (A) transcripts, (B) proteins and (C) PMs. (D, F) Simulated protein profiles from RNAs peaking at (D) ZT0 or (F) ZT16, with (red line) or without light-regulated translation (black line). (E) predicted distribution of protein peak times, with light-regulated translation. Examples of genes with (G) high-amplitude and similar protein (solid line) and PM profiles (coloured lines), or (H) PM profiles that differ from the protein profile. (G, H) protein and PM, left axis; RNA profile (dashed line), right axis. Error bars, S.E. Light/dark indicated by white/black bars.

The regulation was strikingly reversed in the proteome (Fig. 1C, 1D), where the major separation (in PC1) was between samples from light and dark intervals. The early day (ZT4), when translation and chlorophyll biosynthesis GO terms were enriched, was separated from all other timepoints, most strongly from mid-night (ZT16 and 20). There was less separation (in PC2) of the late night (ZT0), when proteins involved in the TCA cycle and transport processes are prominent, from the late day (ZT8-12), when translation and chlorophyll biosynthesis were still enriched. In contrast, the ZT0 timepoint stood out in the phosphoproteome (Fig. 1E, 1F), when PMs enriched for transcription, glucose metabolism, K^+^ and protein transport and ubiquitin-dependent proteolysis functions were clearly separated from the late-day timepoints (ZT8, ZT12). PC2 separated mid-day timepoints (ZT4, ZT8, with enrichment for regulation of gene expression, translation and transmembrane transport) from mid-night (ZT16, ZT20, when mitosis and Ca^2+^ transmembrane transport terms were enriched).

The even distribution of changing RNAs across all the transcriptomic timepoints was not reflected either in the proteome or the phosphoproteome data, where the early day (ZT4) or pre-dawn (ZT0) timepoints, respectively, stood out in the PC analysis. Hierarchical clustering grouped the protein and PM abundance profiles into 8 clusters (termed P1–P8 and PM1– PM8, respectively; Supplementary Figure S2C, S2D), which are coloured on the PC plots in Fig. 1F-1I. GO term enrichment for RNAs, proteins and PMs in the principal component, clustering and peak time analyses is presented in Supplementary Tables S3-S5, with a summary for proteins and PMs in Supplementary Figure S5. We analysed the distribution of individual peak times (Figure 3) to understand these patterns, starting with the proteins.

**Figure 3.**
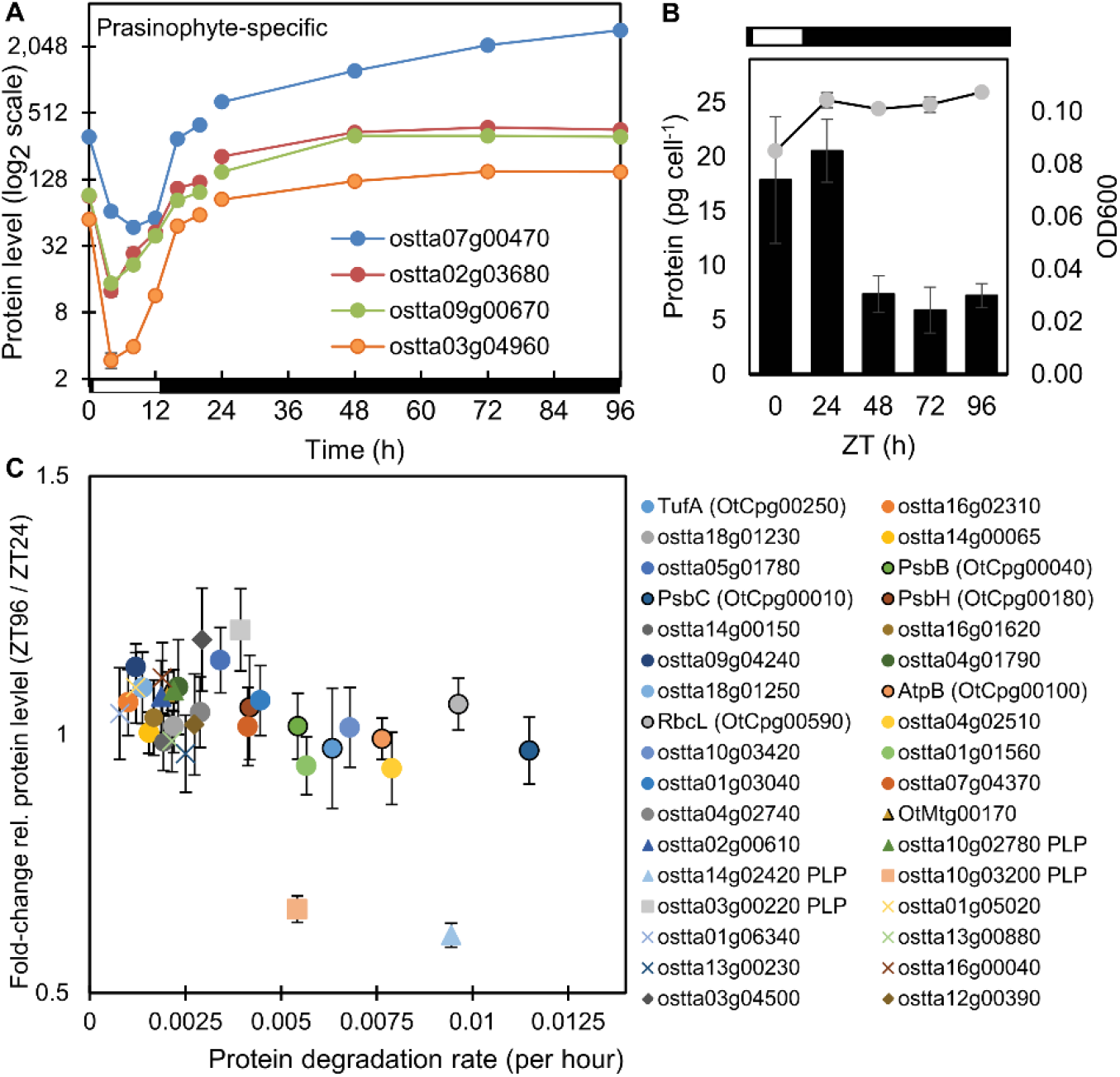
Regulation of dark-accumulating proteins. Protein abundance profiles (A) of rhythmic prasinophyte-specific proteins in cluster P6 in LD and DA conditions. (B) Optical density (OD600; line, right axis) and total protein per cell (columns, left axis) under LD and DA conditions. (C) Correlation of protein degradation rates (Martin *et al*., 2012) and relative protein levels after DA; chloroplast proteins (circles, chloroplast-encoded have solid outline); mitochondrial proteins (triangles, mitochondria-encoded outlined); PLP-enzymes (squares, marked in legend); prasinophyte-specific proteins (diamonds).

### Daytime peaks of protein abundance

Hundreds of transcripts reach peak abundance at every timepoint around the day/night cycle (Fig. 3A) (Monnier *et al*., 2010). In contrast, most protein profiles peaked in the light interval (85% at ZT4-12; Fig. 2B), separating the day and night samples in line with the PC analysis. Metabolic labelling of *O. tauri* has shown ∼5-fold higher protein synthesis rates in the day compared to the night (Martin *et al*., 2012). Consistent with this, our analyses showed translation-related proteins were enriched among the rhythmic proteins with high abundance in the daytime, in PC1, protein cluster P1 and in profiles with daytime peak phase (Supplementary Tables S3-S5, Supplementary Figures S4, S5). We therefore tested whether this light-regulated synthesis alone could explain the observed distribution of protein peaks.

We simulated protein dynamics (Fig. 2D-2F; Supplementary Figure S6) using measured protein synthesis and degradation rates (Martin *et al*., 2012), and an even temporal distribution of rhythmic mRNAs. The simulated distribution of protein profiles matched well with our experimental results (Fig. 2E; Supplementary Figures S6D-S6G), with a slightly stronger daytime preference than in the data. ostta03g04520 is an example of an RNA that peaks at ZT0 and its protein profile (Fig. 2G) was very similar to the predicted protein from such an RNA (Fig. 2D). The overall distribution of protein profiles substantially reflects the light-stimulated translation rate of this organism (see Discussion).

### Unusual, night-time proteins suggest a ‘dark state’

An intriguing pattern of protein regulation stood out from the common, daytime abundance. Protein cluster P6 (and P8) included the rare protein profiles that fell at ZT4 (Supplementary Figure S2C), associated with oxidative metabolism and protein transport GO terms (Supplementary Table S4). Four un-annotated, prasinophyte-specific proteins in cluster P6 not only peaked at night, but were also among the 11, highest-amplitude profiles of all the rhythmic proteins (Fig. 3A). Their dramatic fall in abundance at ZT4 suggested a destabilisation by light, so we tested whether such proteins would remain stable during several days of dark-adaptation (DA).

*O. tauri* cells are photo-autotrophic. Their division is entrained by the LD cycle (Farinas *et al*., 2006) and they arrest transcription in prolonged darkness, when they can survive without growth or division (O’Neill *et al*., 2011). Cell density (optical density at 600nm) in our cultures increased by ∼25% after one LD cycle. Cellular protein content was consistent (18-20 pg cell^-1^) in replicate measures at ZT0 and ZT24 (Fig. 3B). In cultures transferred to three further days of darkness, optical density remained constant but protein content per cell dropped by over 60% on the first day (ZT24 to ZT48) and was then stable to ZT96. This result was suggestive of an altered, but potentially stable, cellular ‘dark state’, which we tested in a further, proteomic timeseries, sampling in darkness at ZT24, 48, 72 and 96.

The proteomic landscape changed less during dark adaptation (DA) than under a standard LD cycle. 98 of the 865 proteins quantified by LC-MS changed levels more than the average and only 64 (7%) also changed more than 1.5-fold (Supplementary Table S6). The 35 significantly-increasing proteins in DA included five transmembrane transporters, a Lon-related protease and two superoxide dismutases, suggestive of nutrient acquisition, protein mobilisation and oxidative stress responses. The four prasinophyte-specific proteins noted above were among the ten most-increasing proteins in DA, confirming their unusual regulation and suggesting a shared function in both standard night-time and the putative ‘dark state’. The most-decreasing among 63 significantly-decreasing proteins in DA was a starch synthase (ostta06g02940). Its abundance declined in the night under LD cycles, as did all 10 of the DA-decreasing proteins that were also rhythmic in LD. The largest functional group of depleted proteins comprised 22 cytosolic ribosomal proteins and translation factors (Supplementary Table S6), suggesting that *O. tauri* selectively mobilised this protein pool in darkness.

The night-abundant, prasinophyte proteins that accumulated in DA, and night-depleted proteins that fell in DA (such as ostta06g02940, above; or PPDK ostta02g04360, Supplementary Figure S7C), suggested that prolonged darkness preserved a night-like state. An alternative explanation was that protein stability in general was altered in the putative dark state. We sought to test that notion, using the protein degradation rates that were previously measured by metabolic labelling in LD conditions (Martin *et al*., 2012). Falling protein abundance under DA was significantly correlated with higher degradation rates in LD (Fig. 3C; R = -0.48, p=0.004, n=34), even among these abundant, stable proteins. We also tested RNA abundance for a subset of these proteins in DA by qRT-PCR, showing stable levels after one day of prolonged darkness (ZT48; Supplementary Figure S8A). The lack of RNA regulation seemed consistent with the lack of transcription in these conditions (O’Neill *et al*., 2011). For example, a further prasinophyte-specific protein ostta03g4500 with a stable RNA level and slightly-increasing protein level in DA also had among the lowest protein degradation rates in LD (Fig. 3C), and was among the most-detected proteins in these conditions (Supplementary Figures S2A, S8B). The RNA data and protein degradation rates suggested that the prasinophyte-specific proteins accumulated due to a focussed, regulatory mechanism, rather than generalised refactoring of the proteome (see Discussion).

### A phospho-dawn of protein modification

In contrast to the many daytime-peaking protein profiles, 39% of the changing phosphopeptide motifs (PMs) peaked in abundance at ZT0 (Fig. 2C), double the proportion of any other timepoint. The ZT0 samples were harvested before lights-on, so this ‘phospho-dawn’ anticipated the transition and was not due to light-stimulated translation. Other, high-amplitude PM profiles tracked the levels of their cognate proteins, with little evidence of regulated phosphorylation (Fig. 2G). We therefore tested the contribution of protein levels to PM profiles more broadly, among the 138 genes that were quantified in both protein and PM datasets (Supplementary Figures S7A-B). This subset of 261 protein-PM pairings included proteins peaking at all timepoints, and PM profiles that reflected the peak time distribution of the full dataset. 80% of the PMs peaked at different timepoints than their cognate protein (Supplementary Figure S7C; examples in Fig. 2H). The LHC linker protein CP29 (ostta01g04940) illustrates one pattern: its protein level rises in the light while a PM is de-phosphorylated (Supplementary Figure S7C) adjacent to a target site of chloroplast kinase STN7 in Arabidopsis (Schönberg *et al*., 2017).

To test the phospho-dawn pattern by a different method, we estimated the bulk protein phosphorylation across the diel cycle using protein gel staining (Supplementary Figures S9A-B). The proportion of phosphorylated proteins was lowest in the daytime and increased during the night to peak at ZT0 (Supplementary Figures S9C). Total phosphorylation was therefore broadly consistent with the distribution of PM profiles (Fig. 2C). Taken together, these results indicate that a regulator other than light or protein abundance controls the *O. tauri* phosphoproteome before dawn. Below, we report phosphosite sequences that suggested its identity.

### Functions of proteins with rhythmic phospho-motifs

The LD datasets confirmed that protein phosphorylation profiles often diverged from protein abundance. Colour-coding in Fig. 1H shows that clustering of the phospho-motif (PM) profiles aligned with the PC analysis more clearly than for the lower-amplitude, protein profiles (Fig. 1F). The largest cluster PM1 reflected the profiles that peaked in the ZT0 timepoint, which PC analysis also highlighted (Supporting Figure S2D). Phosphopeptide enrichment allowed the detection of many regulatory proteins, including PMs on predicted CONSTANS-like B-box transcription factors (OtCOL) related to the plant clock protein TOC1 (Fig. 4), and on the RWP-RK mating-type factor ostta02g04300 (Blanc-Mathieu *et al*., 2017). PM1 also includes the predicted CK2 target site pS10 in the clock protein CCA1 (ostta06g02340; Fig. 4), close to the homologous location of a CK2 site in Arabidopsis CCA1 (Lu *et al*., 2011).

**Figure 4.**
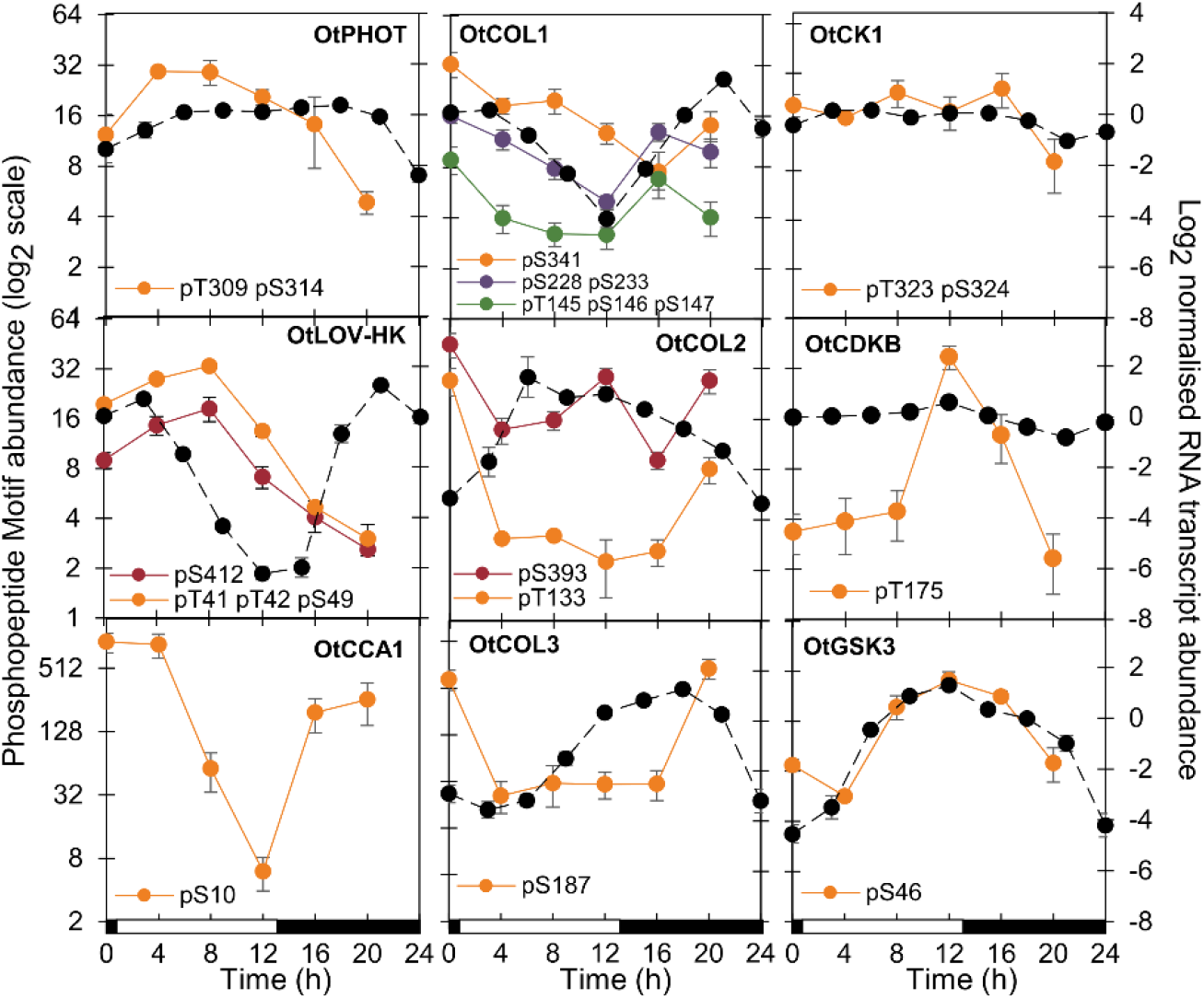
Protein and phosphopeptide motif regulation. Phosphomotif (coloured lines) and RNA profiles (Monnier *et al*., 2010)(dashed lines) of the photoreceptors, clock components, transcription factors and kinases indicated, under LD. Left axis range 2^6^ (64-fold) except OtCCA1 (PM changes 150-fold) and OtCOL2 (PMs change up to 20-fold). Right (RNA) axis range 12, for log2 data (2^12^=4096-fold in untransformed data). Error bars, S.E. Light/dark indicated by white/black bars. PHOT, phototropin photoreceptor; LOV-HK, LOV domain – histidine kinase photoreceptor; COL, CONSTANS-like transcription factor.

PMs in cluster PM3 peaked in the light, consistent with many protein profiles (examples in Fig. 2G). PMs on the photoreceptors phototropin and LOV-HK illustrate these daytime profiles (Fig. 4). Protein functions predicted to regulate transcription, metal ion transport and protein phosphorylation are enriched in this cluster (summarised in Supplementary Figure S2F; Supplementary Table S4) and in profiles with daytime peaks (Supplementary Figure S4B; Supplementary Table S5).

In contrast, the PM2, PM4, PM7 and PM8 clusters peaked at ZT16, with or without accumulation in daytime (Supplementary Figure S2D). These clusters are enriched for PMs on protein kinases including cell-cycle-related kinases (Supplementary Figures S2F, S4B; Supplementary Tables S4 and S5). We therefore analysed the phospho-regulators that might control these profiles, including potential contributions to non-transcriptional timing.

### Phase-specific target sites

We first analysed motifs of amino acids that were enriched in rhythmic PMs, compared with all quantified phosphopeptides to avoid potential detection bias due to PM abundance. PMs that peaked at ZT16 were strikingly enriched for the proline-directed motif [pS/pT]P (Fig. 5B-C). This strongly implicates the CMGC family of protein kinases, including Cyclin-Dependent Kinases (CDKs) and GSK. Consistent with this, the profiles of PMs with predicted GSK target sequences also most often peaked at ZT16 (Supplementary Figure S10A-B). Levels of *GSK3* RNA and a PM on GSK3 peaked at ZT12 (Fig. 4), though the auto-phosphorylation site pY210 was not rhythmic (Supplementary Fig. S2B; Supplementary Table S2). More specific CDK target motifs [pS/pT]PXX[K/R] were enriched at ZT12, consistent with the known timing of cell division (Farinas *et al*., 2006; Moulager *et al*., 2007) and the peak level of the activation phospho-site of CDKB (Fig. 4). During the day (ZT4 and 8), enrichment of hydrophobic residues at positions -5 and +4 is suggestive of the SnRK consensus (Vlad *et al*., 2008), the plant kinase most related to animal AMPK.

**Figure 5.**
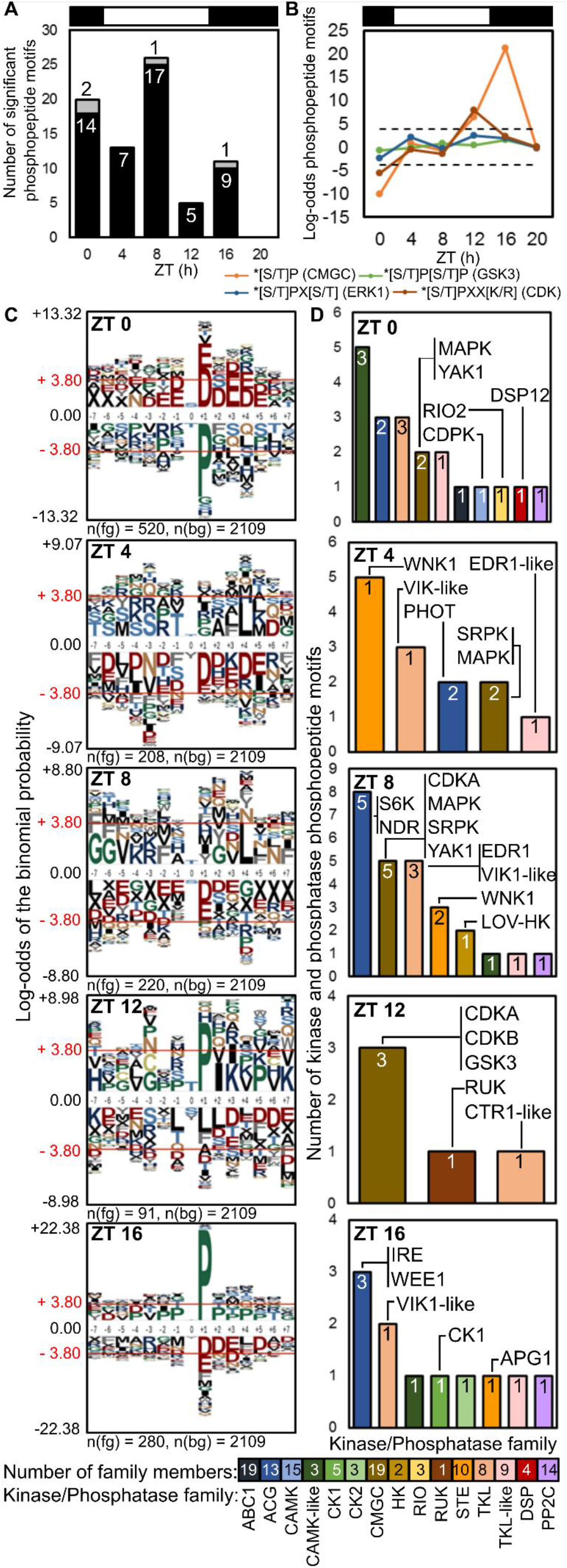
Motif enrichment and rhythmic protein kinases and phosphatases under LD. (A) Rhythmic PMs peaking at each timepoint on protein kinase (black) and phosphatase (grey) proteins (numbers). (B) Enrichment of proline-directed motifs, for kinases shown in the legend (dashed line, *p*-value = 0.05). (C) pLogo sequence motifs of rhythmic PMs peaking at each timepoint (foreground; fg), relative to all detected phosphopeptides (background; bg). ± 3.80 indicates *p*-value = 0.05, residues above and below axis are over- and under-represented, respectively. (D) Rhythmic PMs by kinase/phosphatase family, annotated with example proteins.

In contrast, acid([D/E])-directed target motifs were significantly enriched among the many rhythmic PMs that peaked at ZT0 and the proline-directed motifs were depleted (Fig. 5C). Conversely, these acid-directed motifs were depleted on PMs peaking at ZT16 or ZT4, suggesting a strong phase-specificity. Considering the more specific, predicted target sites for the clock-related protein kinases (Supplementary Figures S10A), predicted CK1 targets were most abundant, and most often peaked at ZT0. Predicted CK2 target sequences were even more phase-specific, with at least 5-fold more peaking at ZT0 than at other times. Thus predicted targets of the clock-related kinases CK1 and CK2 both contribute to the phospho-dawn profiles, in antiphase to the evening peaks of proline-directed phospho-sites.

### Rhythmic regulation of the kinome

The protein abundance of the three detected protein kinases and two phosphatases was not rhythmic (Supplementary Table S1). We therefore analysed the 68 rhythmic PMs on protein kinases and five PMs on protein phosphatases, as candidate mediators of rhythmic phosphorylation (Figs. 5A, 5D). The PMs on kinases represent 8% of the total, though protein kinase genes comprise ∼1.5% of the genome. Indeed, the most heavily-phosphorylated protein with 14 PMs was the WITH NO LYSINE (WNK) kinase that might target clock proteins in Arabidopsis (Murakami-Kojima *et al*., 2002) (Supplementary Table S2; Supplementary Figure S10C). The most-changing PM on a predicted protein phosphatase was pT175 in ostta11g02830, related to human Dual-specificity phosphatase DUSP12 (Fig. 5D).

Among the clock-related protein kinases, we note the dusk-peaking PM of GSK3 (above). CK2 subunits were not detected in our data and the PM on CK1 was not strongly rhythmic (Fig. 4). 21 other protein kinases bore rhythmic PMs that are predicted targets of these clock-related kinases (Supplementary Figures S10C).

Around mitosis at ZT12-16, significantly peaking PMs were detected on cell cycle regulators CDKA, CDKB and WEE1 (Fig. 5D). Kinase PMs peaking at ZT4-8 included Serine-Arginine Protein Kinases (SRPKs), MAPKs, CDKA and a site on Yet Another Kinase (YAK1). PMs that peaked at ZT0, coincident with the phospho-dawn, included RIO2, YAK1 and CDPK, all implicated in cell cycle regulation and progression (Garrett *et al*., 1991; LaRonde-LeBlanc and Wlodawer, 2005). RIO’s are among the few kinase families shared with the Archaea (Kennelly, 2014), making them candidate contributors to an ancient, non-transcriptional oscillator (Edgar *et al*., 2012).

## Discussion

### The diel proteome and phosphoproteome

Our results contribute to understand the ‘reactive’ and ‘anticipatory’ components of protein regulation in the green lineage under diel (LD) cycles (Mehta *et al*., 2021). A small fraction of the *O. tauri* proteins quantified here were rhythmic (just under 10%), compared to a majority (58%) of the phosphomotifs (PMs). Most protein profiles peaked in daytime, consistent with the ‘reactive’ effect predicted from the light-regulated translation that was previously measured in this organism (Martin *et al*., 2012), and with enrichment of translation-related functions among daytime-peaking proteins. This result reinforces the dangers of using RNA profiles as a proxy for biological function. It further supports our prediction that “translational coincidence” should alter the *O. tauri* proteome in different day lengths, as some rhythmic RNAs will coincide with light-stimulated translation only in long days (Seaton *et al*., 2018).

In contrast, the largest number of PM profiles peaked in the pre-dawn, ZT0 timepoint. This anticipatory ‘phospho-dawn’ might be controlled by the circadian clock. Circadian regulation would be expected to persist under constant conditions, which were not tested here. Studies in Arabidopsis under constant light, however, identified a high fraction of rhythmic phosphopeptides that peaked at subjective dawn (Choudhary *et al*., 2015)(Krahmer *et al*., 2018), suggesting a similar, circadian-regulated phospho-dawn in higher plants. Such phospho-regulation might prepare green cells for daytime functions and/or end night-time activities, before light-stimulated translation facilitates new protein synthesis.

Acid-directed target sites were clearly enriched at ZT0, implicating the clock-related kinases CK1 and CK2 in regulating the phospho-dawn in *O. tauri*. Enrichment of proline-directed target sites occurs in antiphase, at ZT12-16, which implicates the 19 CMGC-class kinase proteins (Hindle *et al*., 2014) including CDKs, MAPKs and GSK3. These phase-specific enrichments were clearer than in the Arabidopsis studies, suggesting that the minimal kinase-target network of *O. tauri* might be easier to resolve in future. Comparison to the rhythmic phosphoproteome in animals is limited, because the most-rhythmic kinase Akt (also known as Protein Kinase B) in mouse liver (Robles *et al*., 2016) is absent from the green lineage (Hindle *et al*., 2014).

The low overall rhythmicity (<10%) in the partial proteome quantified here is consistent with similar studies in Arabidopsis, which identified 0.1-1.5% rhythmic proteins from 7-9 % of the proteome in LD, using iTRAQ labelling with similar statistical criteria to ours (Baerenfaller *et al*., 2015; Baerenfaller *et al*., 2012), or 4-7% rhythmic proteins from 4% of the proteome under constant light using a gel-based approach (Choudhary *et al*., 2016). Our results provide 11% coverage in the minimal *O. tauri* proteome, with a simpler experimental protocol. Broader coverage of this proteome was reported after our preprint was released (Kay *et al*., 2021), in experiments that included a depletion of abundant proteins, among several technical differences. Their higher reported fraction of rhythmic proteins might reflect the detection of low-abundance proteins and/or analysis with no minimum amplitude threshold.

### The ‘dark state’ is indirectly associated with lipid synthesis

Among the rhythmic proteins reported here, some of the most highly-regulated were four prasinophyte-specific sequences (unnamed proteins ostta02g03680, ostta03g04960, ostta07g00470, ostta09g00670; Figure 3A) along with ostta03g04500 (Supplementary Figure S2A). These proteins accumulated further in prolonged darkness (Figure 3A). We previously showed that *O. tauri* stop transcription and cell division in those conditions. Cultures resume gene expression and growth upon return to LD cycles, suggesting that dark adaptation induces a state of cellular quiescence. The ecological relevance of a quiescent ‘dark state’ for photo-autotrophic, surface-dwelling *O. tauri* is not immediately obvious. However, *Ostreococcus* relatives can persist under the Polar Night (Joli *et al*., 2017), and quiescent forms in other phytoplankton (Roy *et al.,* 2014) can be ecologically important in benthic-pelagic coupling (Marcus and Boero, 1998). Cells near the deep chlorophyll maximum (Cardol *et al*., 2008) could be moved into the dark, benthic zone by turbulence, to return later *via* upwelling (Collado-Fabbri *et al*.,2011; Countway and Caron, 2006). Understanding the laboratory ‘dark state’ might therefore have ecological relevance.

Protein content dropped significantly between 12h and 36h of darkness (ZT24 to ZT48) but was then stable. Proteins associated with cytosolic translation were notably depleted (Supplementary Table 6), rather than abundant, chloroplast proteins involved in photosynthesis. Photosynthetic functions might be particularly important to recover from quiescence, similar to the rapid regrowth observed after nutrient starvation (Liefer *et al.,* 2018).

A preprint coincident with our first report showed that three of these night-expressed, prasinophyte-specific proteins accumulated strongly in *O. tauri* when the growth medium was depleted of nitrogen, under LD cycles (Smallwood *et al*., 2018a). The most-depleted protein in their conditions was the same starch synthase (ostta06g02940) that fell most in abundance under our prolonged dark treatment. Nitrogen depletion is commonly used to induce lipid synthesis in algae, in the context of third-generation biofuel production (Zienkiewicz *et al*., 2016). Prolonged darkness and/or hypoxia can also induce lipid accumulation, and hypoxia can occur in dark-adapting algal cultures due to continued respiration (Hemschemeier *et al*., 2013). Our culture conditions included sorbitol and glycerol in the growth medium (O’Neill *et al*., 2011), which are required for viability in prolonged darkness. Our ‘dark state’ proteome might therefore reflect active lipid synthesis from these substrates.

*O. tauri* can form both intracellular lipid droplets and extracellular droplets in membrane-bound ‘pea-pod’ structures (Smallwood *et al*., 2018*b*). Lipid droplets in other algae include major proteins that are restricted to limited taxonomic groups (Zienkiewicz *et al*., 2016). Some of these are predicted to have all-alpha-helical structure, including the Major Lipid Droplet Protein Cre09.g405500 of *Chlamydomonas reinhardtii* or the Lipid Droplet Surface Protein of the diatom *Nannochloropsis oceanica*. Protein structure homology modelling aligned ostta02g03680 with a human BAR domain dimer, an all-helical protein domain that can sense and create membrane curvature (Simunovic *et al*., 2015)(Supplementary Figure S11), suggesting that this *O. tauri* protein might also be involved in lipid droplets. *N. oceanica* lipid synthesis and LDSP accumulation is highly rhythmic but day-phased (Poliner *et al*., 2015). The night-expressed proteins in *O. tauri* indirectly suggest a different regulation of lipid synthesis, that could have biotechnological relevance.

## Abbreviations

PM: phosphopeptide motif;
LD: light-dark cycles;
ZT: Zeitgeber Time;
DA: dark adaptation;
PC: principal component;
CK1: casein kinase 1;
CK2: casein kinase 2;
GSK3: Glycogen Synthase Kinase 3;
CMGC: Cyclin-dependent kinase, Mitogen-activated protein kinase, Glycogen synthase kinase, CDC-like kinase;
CCA1: Circadian Clock Associated 1 protein.

## Supplementary Data Summary

**Supplementary Figure S1. Identification of outlier phosphopeptide replicate 4E**.

**Supplementary Figure S2. Most-detected protein and PM profiles**.

**Supplementary Figure S3. Changing PMs on non-changing proteins**.

**Supplementary Figure S4. Clustered protein and PM profiles with enriched functions**.

**Supplementary Figure S5. Phase-specific GO term enrichment**.

**Supplementary Figure S6. Simulation of light-regulated translation**.

**Supplementary Figure S7. Loci identified in both LD protein and phosphopeptide motif datasets**.

**Supplementary Figure S8. Regulation of proteins tested under Dark Adaptation (DA)**.

**Supplementary Figure S9. Protein and phospho-protein abundance in LD cycle**.

**Supplementary Figure S10. CK1, CK2 and GSK3 kinase targets and phosphorylation sites in rhythmic kinases**.

**Supplementary Figure S11. Structural homology of rhythmic, prasinophyte-specific protein**.

**Supplementary Table S1. Proteins quantified under LD**.

**Supplementary Table S2. Phosphopeptide Motifs (PMs) quantified under LD**.

**Supplementary Table S3. GO term enrichment among RNA, proteins and PMs contributing to PCA**.

**Supplementary Table S4. GO term enrichment among RNA, proteins and PMs in clusters**. Individually-significant, rhythmic protein profiles are considered, to provide sufficient numbers for enrichment analysis. Only BH-corrected significant PM profiles with >1.5-fold changes are considered.

**Supplementary Table S5. GO term enrichment among rhythmic proteins and PMs by peak/trough times**. Only BH-corrected significant protein or PM profiles with >1.5-fold changes are considered.

**Supplementary Table S6. Proteins quantified under DA**.

## Acknowledgements

We are very grateful to K. Kis, L. Imrie and D. Kelly for expert technical help, to B. Kolody and A. Dodd for helpful discussion, to M. Hirsch-Hoffmann and K. Baerenfaller for support on pep2pro, and to C.R. Smallwood, J.E. Evans and colleagues for clarifying the comparison to their work. The proteomics analyses were carried out by the EdinOmics research facility (RRID: SCR_021838) at the University of Edinburgh. Funded by the Biotechnology and Biological Sciences Research Council (UKRI-BBSRC award BB/J009423/1). For the purpose of open access, the author has applied a Creative Commons Attribution (CC BY) licence to any Author Accepted Manuscript version arising from this submission.

## Author Contributions

Z.B.N. and M.M.H. contributed equally to this study. Investigation and formal analysis, Z.B.N., S.F.M. and T.L.B; formal analysis (bioinformatics), M.M.H.; formal analysis (mathematical modelling), D.D.S.; conceptualisation and methodology, A.J.M., T.I.S., T.L.B., S.F.M., M.M.H. and Z.B.N.; funding acquisition and supervision, A.J.M., T.I.S. and T.L.B.; writing, Z.B.N., M.M.H., T.L.B. and A.J.M.

## Conflicts of Interest

The authors declare no competing financial interests.

## Data Availability

The OTTH95 strain is available from the CCAP (www.ccap.ac.uk) and RCC (roscoff-culture-collection.org) stock centres. Mass spectrometry proteomics data have been deposited in the ProteomeXchange Consortium via the PRIDE partner repository with the dataset identifiers: LD global proteomics, PXD001735; LD phosphoproteomics, PXD001734; DA global proteomics, PXD002909. The LC-MS data were also previously available in pep2pro at www.pep2pro.ethz.ch (Assemblies ‘Ostreococcus tauri Light:dark cycle,LD global’, ‘Ostreococcus tauri Light:dark cycle,LD phospho’, ‘Ostreococcus tauri dark adaptation,DA global’). Processed data lists are provided in the Supplementary Information.

## Supplementary Figure Legends

**Supplementary Figure S1.**
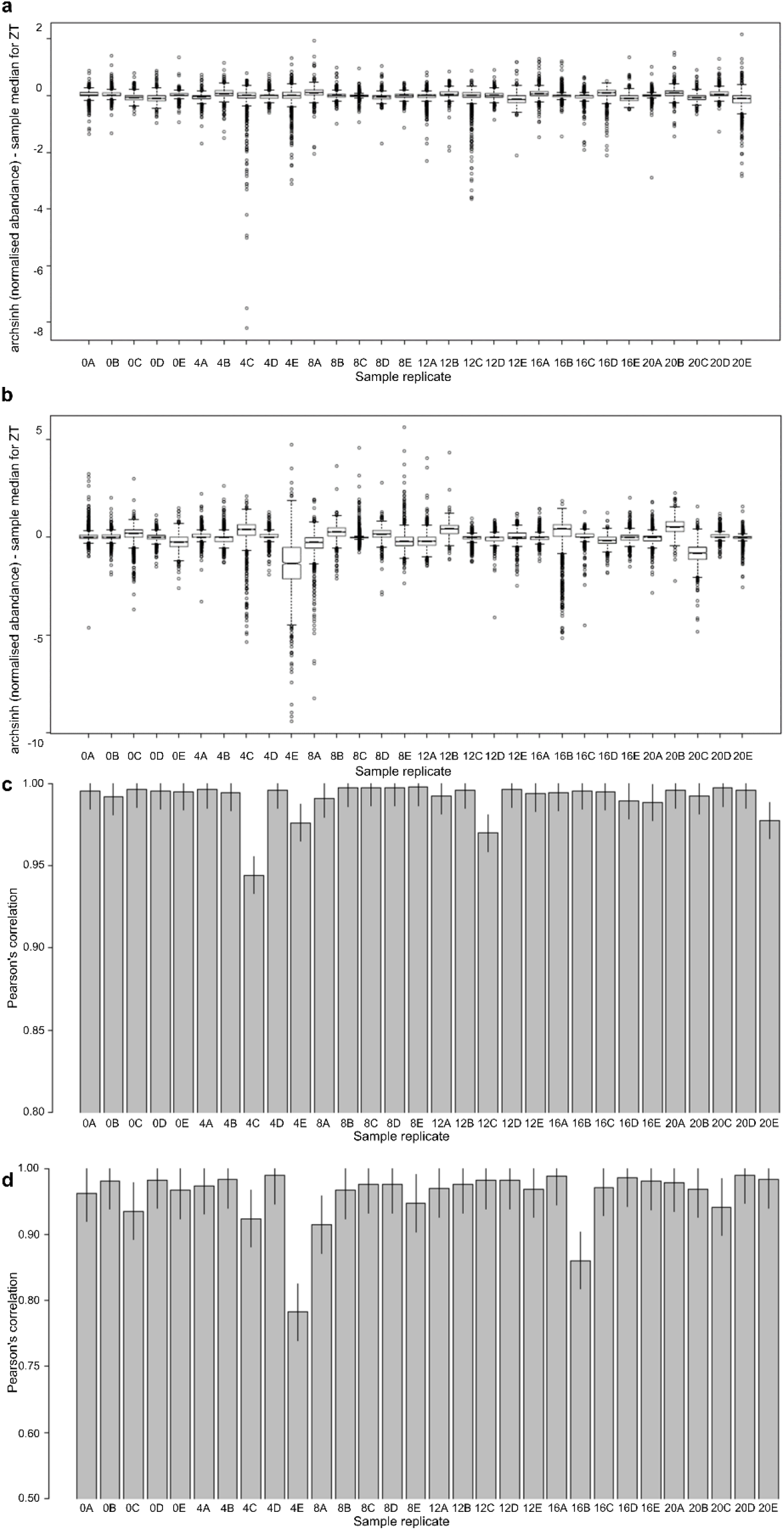
Identification of outlier phosphopeptide replicate 4E. Pearson’s correlation for (a) proteins and (b) phosphopeptide motifs and sample replicate *r*^*2*^ respective to median abundance at a ZT for (c) proteins and (d) phosphopeptide motifs. Note differing scales in (a,b), (c,d).

**Supplementary Figure S2.**
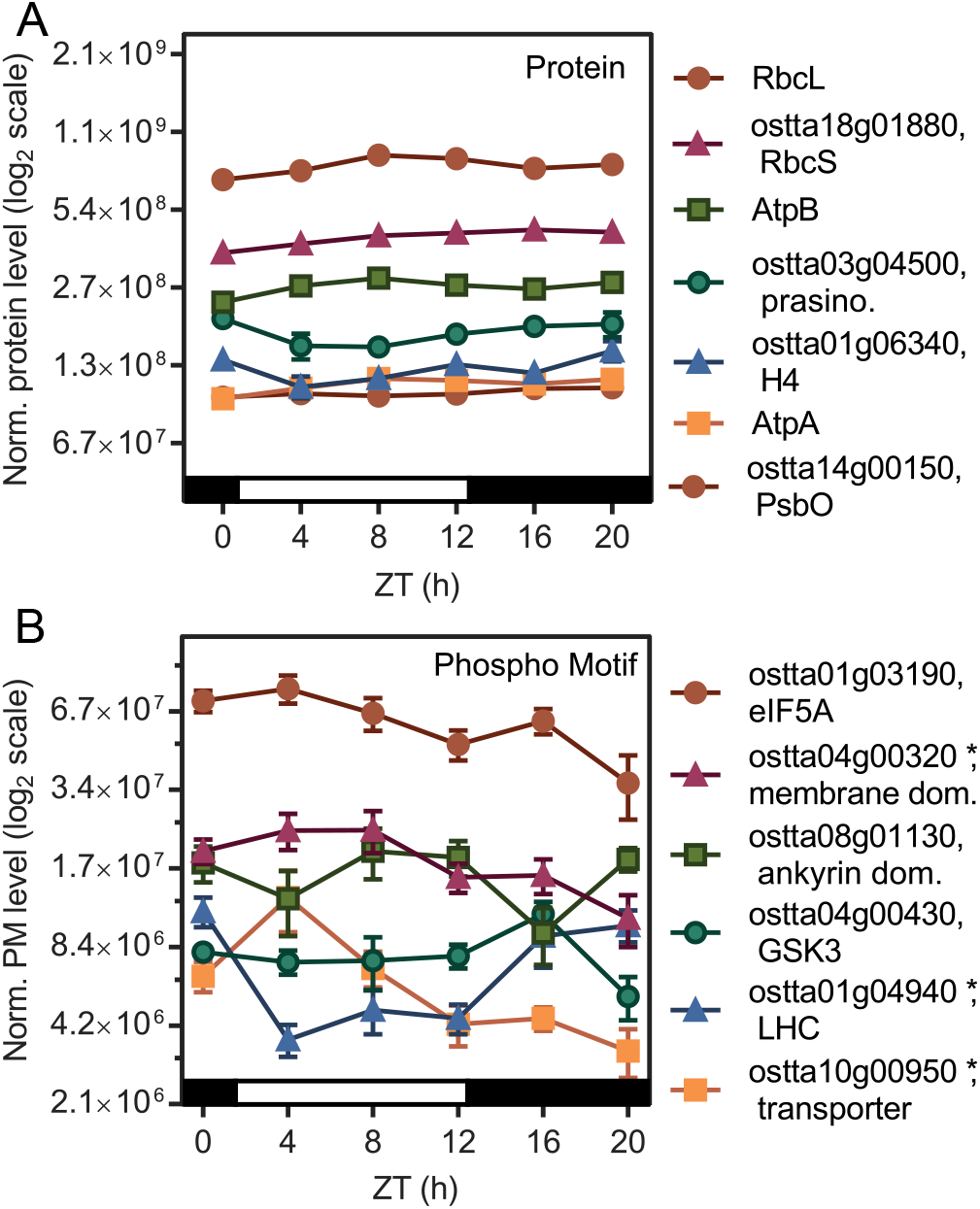
Most-detected protein and PM profiles. with comprehensive heat maps, clusters and enriched functions. Highly-abundant proteins (a) and PMs (b) under LD conditions (* marks rhythmic PMs). Error bars, S.E. Light/dark indicated by white/black bars, above.

**Supplementary Figure S3.**
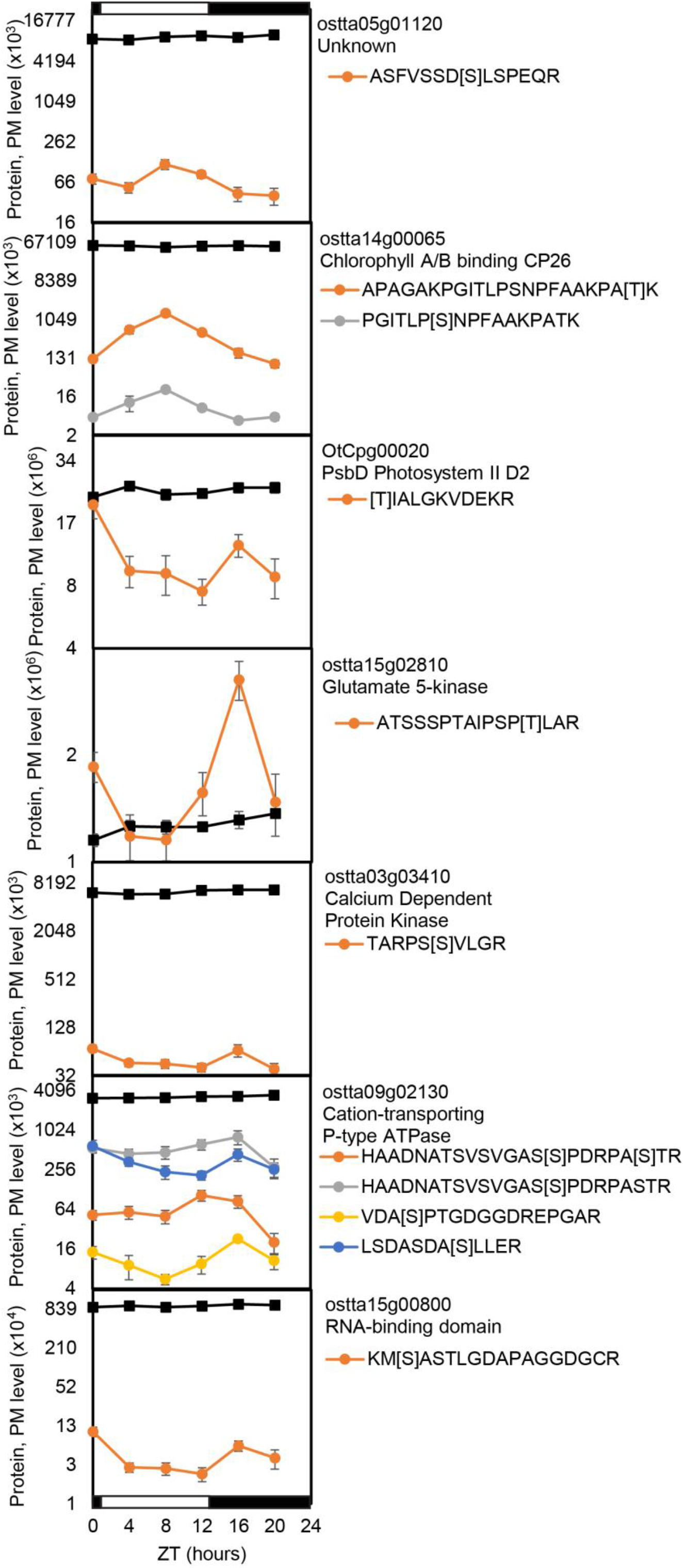
Changing PMs on non-changing proteins. Significantly non-changing proteins (black lines) determined by two one-sided tests (TOST; ε = 0.3), plotted with their rhythmic phosphopeptide motifs ± S.E., square brackets show phosphorylated residue. Light/dark indicated by white/black bars.

**Supplementary Figure S4.**
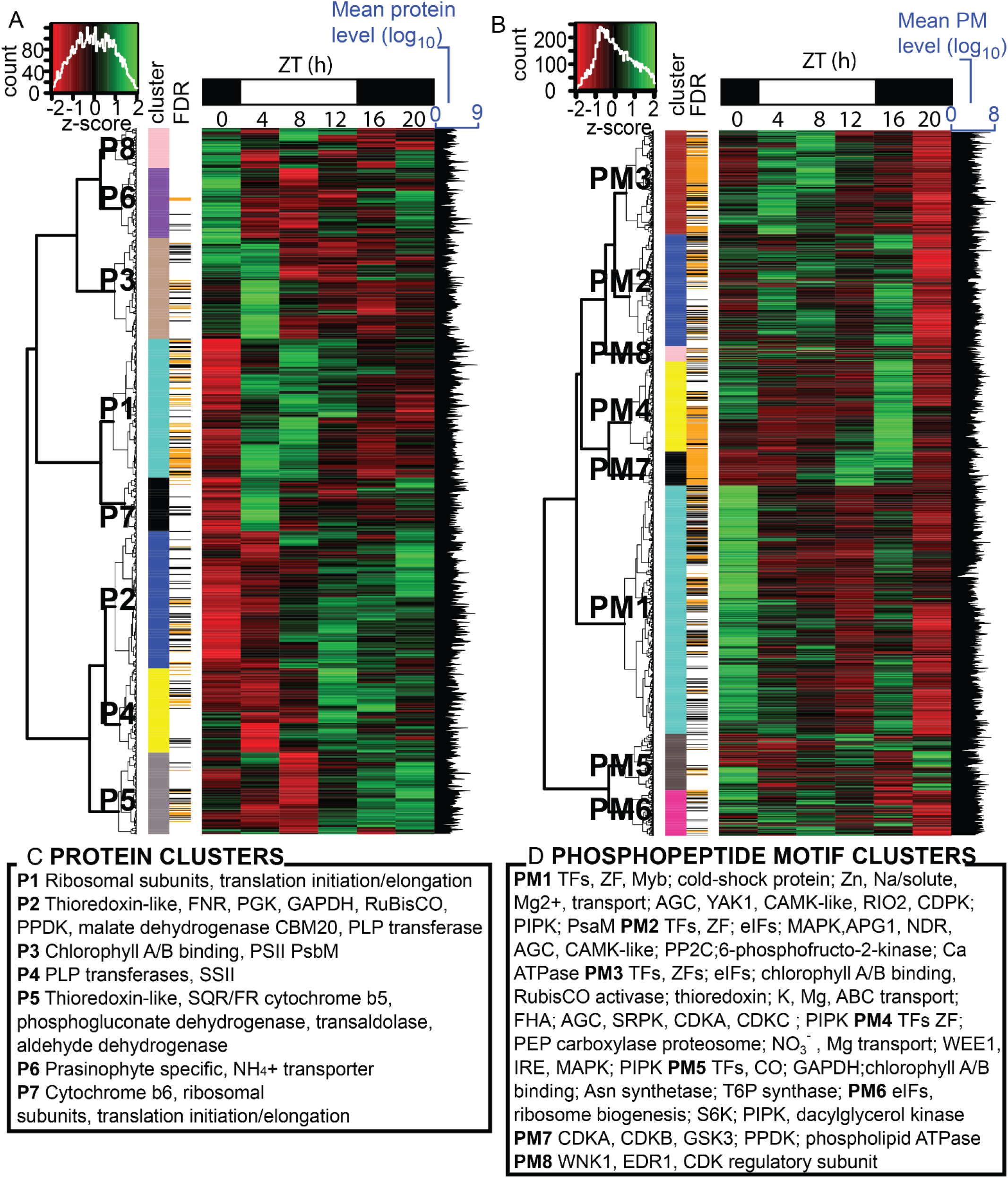
Clustered protein and PM profiles with enriched functions. Heat maps of median-normalised (A) protein and (B) PM abundance, with insets top left showing the distribution of levels and colour scale. Clusters P1-8 or PM1-8 are shown, colours in ‘cluster’ track are as in Fig. 1; FDR track shows >1.5 fold-change and BH FDR adjusted *p*-value <0.05 (black line) or <0.01 (orange line); bars to right of each panel show the mean protein or PM abundance (log_10_ scale). Light/dark indicated by white/black bars, above. (C, D) Examples of significantly-changing proteins and PMs in each cluster (as noted in the main text).

**Supplementary Figure S5.**
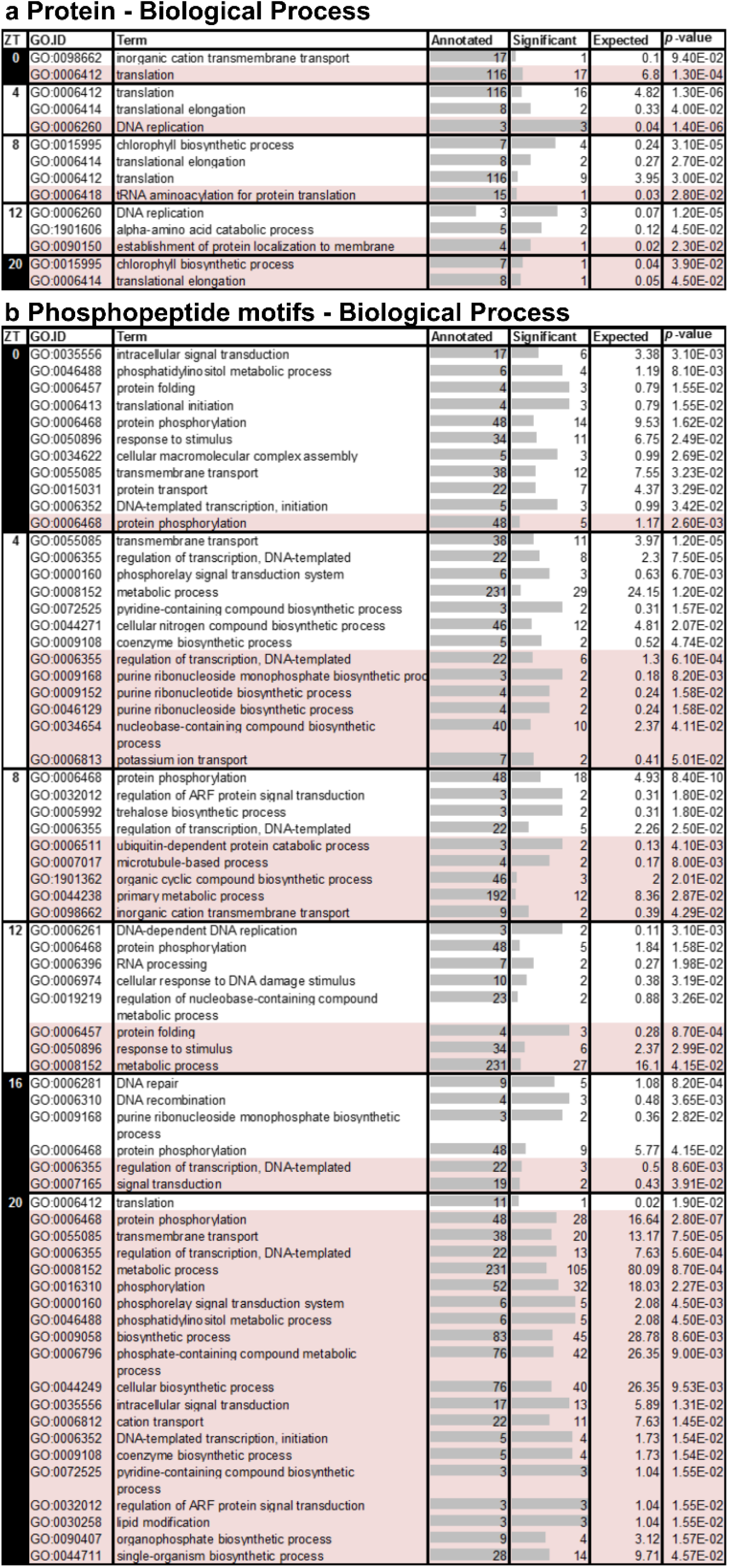
GO enrichments for peaks and troughs. GO Biological Process term enrichment for rhythmic (a) proteins and (b) phosphopeptide motifs, that was significant (Fisher’s exact test p-value <0.05) in profiles with peak (no shading) or trough (pink shading) time at each timepoint. Light/dark indicated by white/black column. Grey bars represent proportion of significant terms identified with respect to total number of background annotated terms.

**Supplementary Figure S6.**
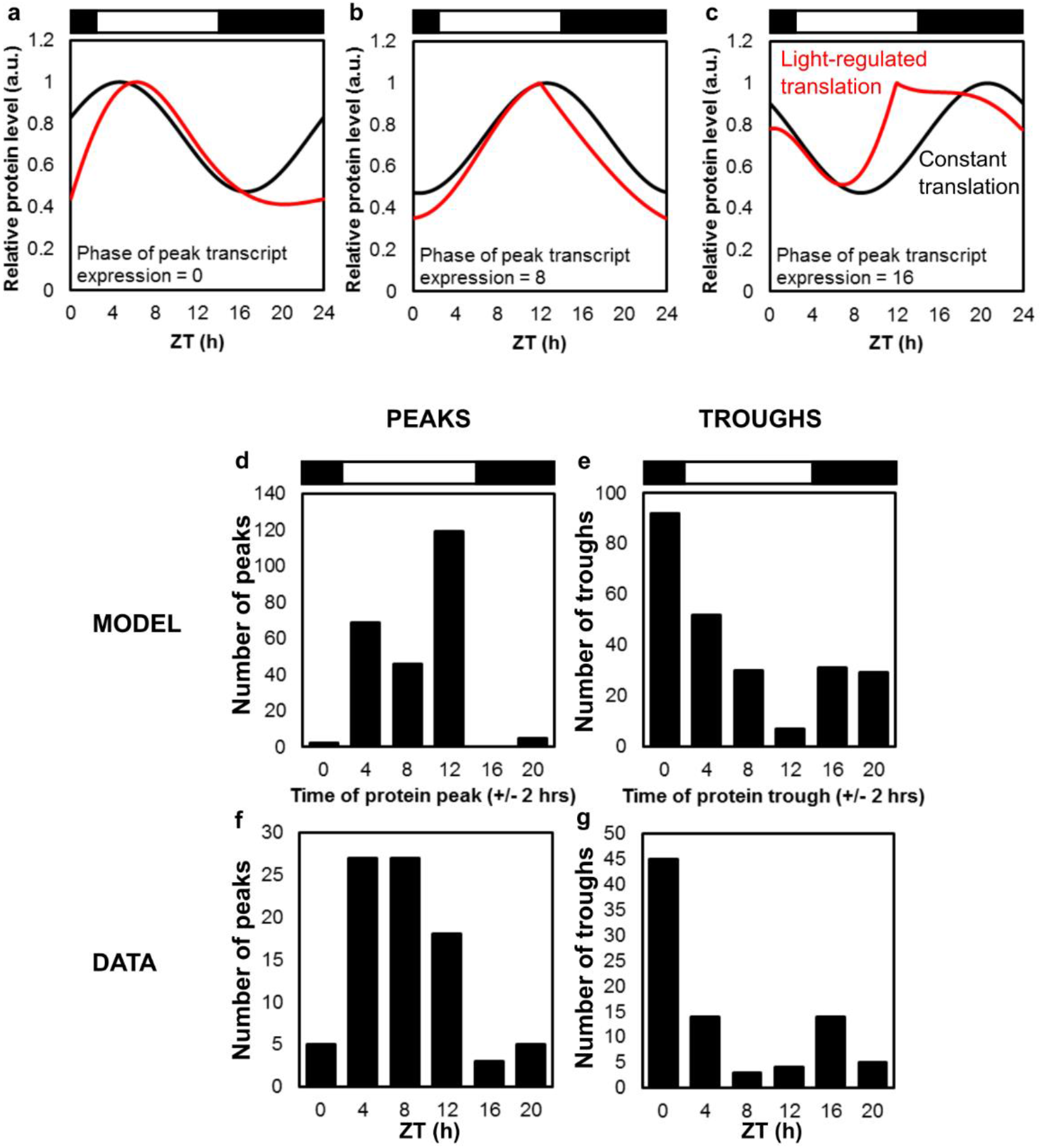
Simulation of light-regulated translation. (a-c) Simulation of protein dynamics for an RNA with peak expression at ZT0 (a), ZT8 (b) and ZT16 (c), with observed, light-regulated translation rate (red lines) or with constant translation rate (black lines). Distribution of protein peaks (d,f) and troughs (e,g) for the model with light-regulated translation (d,e) compared to data (f,g). Distributions for constant translation would reflect the distribution of RNA profiles.

**Supplementary Figure S7.**
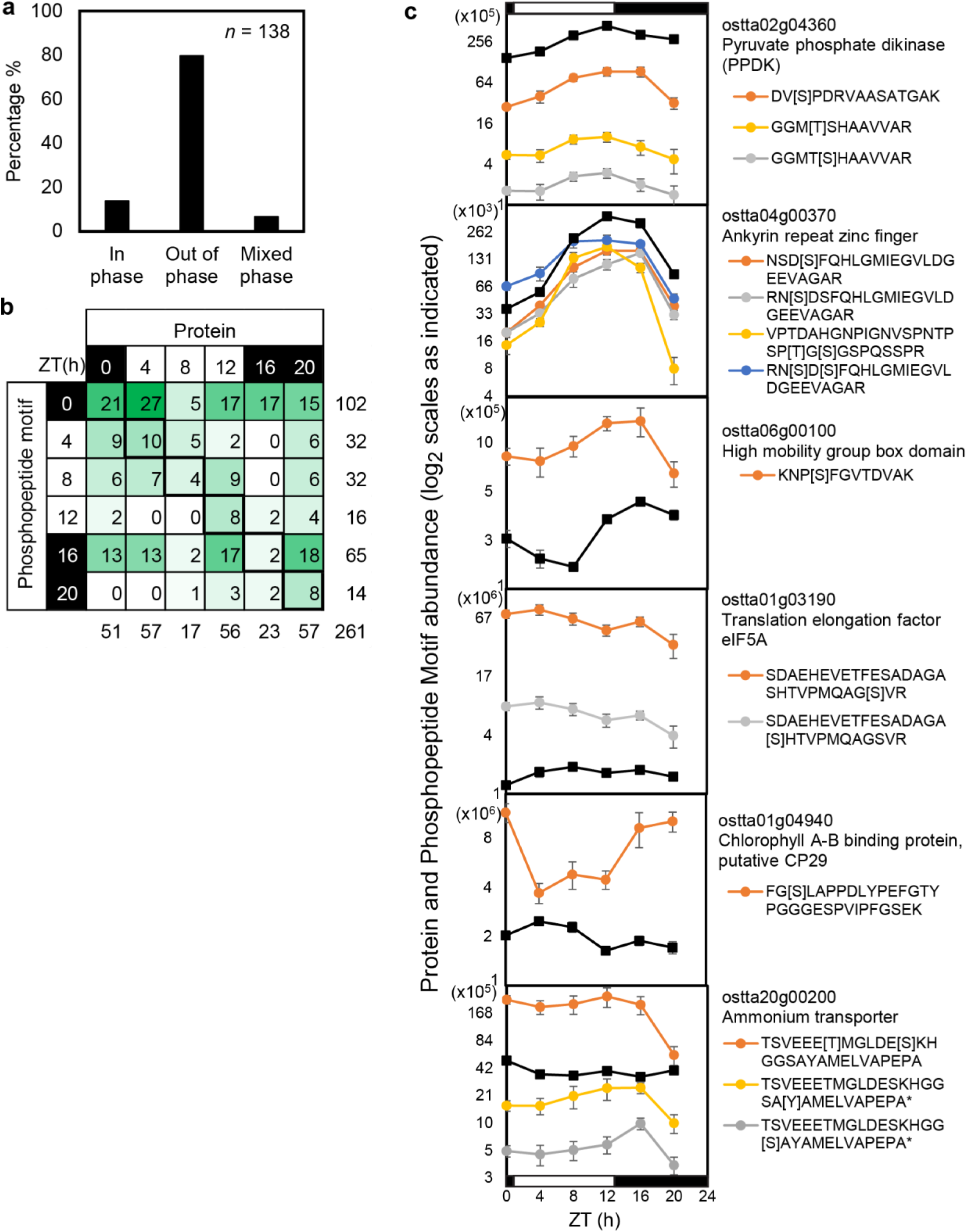
Loci identified in both LD protein and phosphopeptide motif datasets. (a, b) Peak time is compared for genes identified in both LD protein and phosphopeptide motif datasets, with examples (c). (a) Mixed phase: multiple PMs, peaking at same and different times from cognate protein. Green shading in (b) follows number per bin. Plotting conventions in (c) follow Fig. 2c, 2d.

**Supplementary Figure S8.**
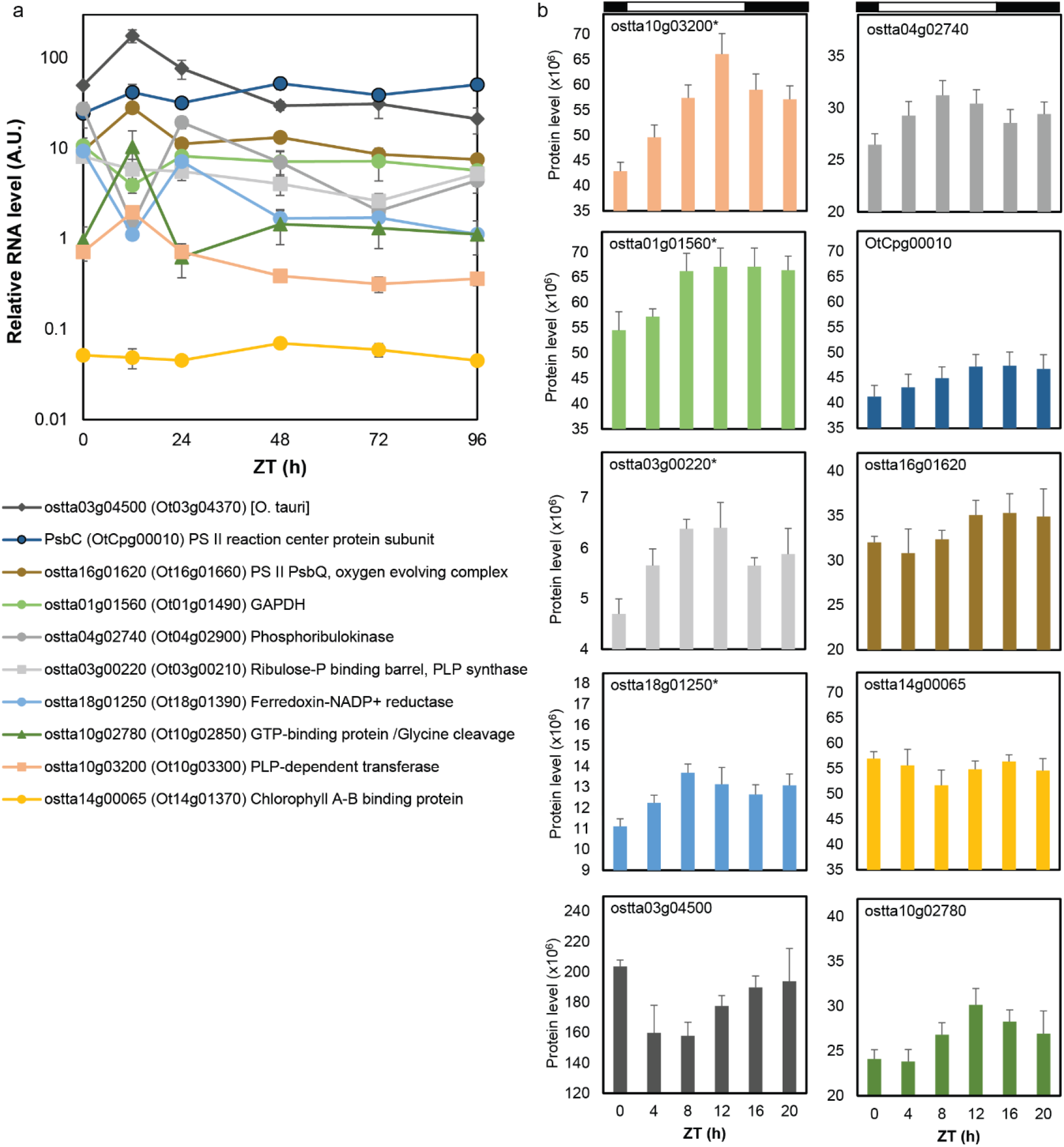
Regulation of proteins tested under Dark Adaptation (DA). For ten proteins compared in the DA and metabolic labelling (Martin *et al*., 2012) data (Fig. 4c), (a) RNA abundance under LD and DA conditions from qRT-PCR assays, and (b) protein profiles under LD. *, rhythmic proteins. Error bar, S.E.

**Supplementary Figure S9.**
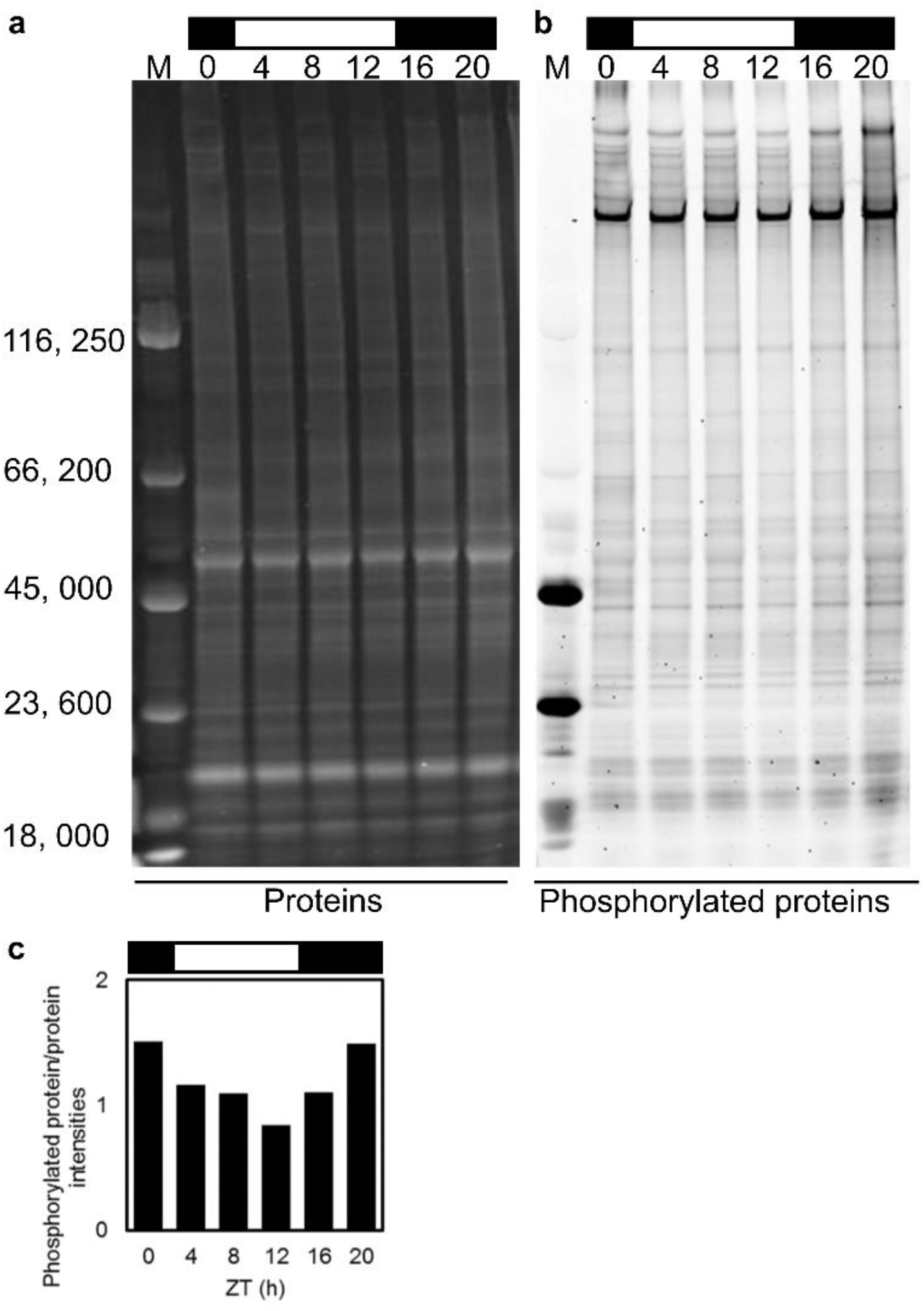
Protein and phospho-protein abundance in LD cycle. Stained gels showing changes in (a) protein and (b) phosphorylated protein abundance in LD, with (c) ratio of quantified, phosphorylated protein to total protein intensity.

**Supplementary Figure S10.**
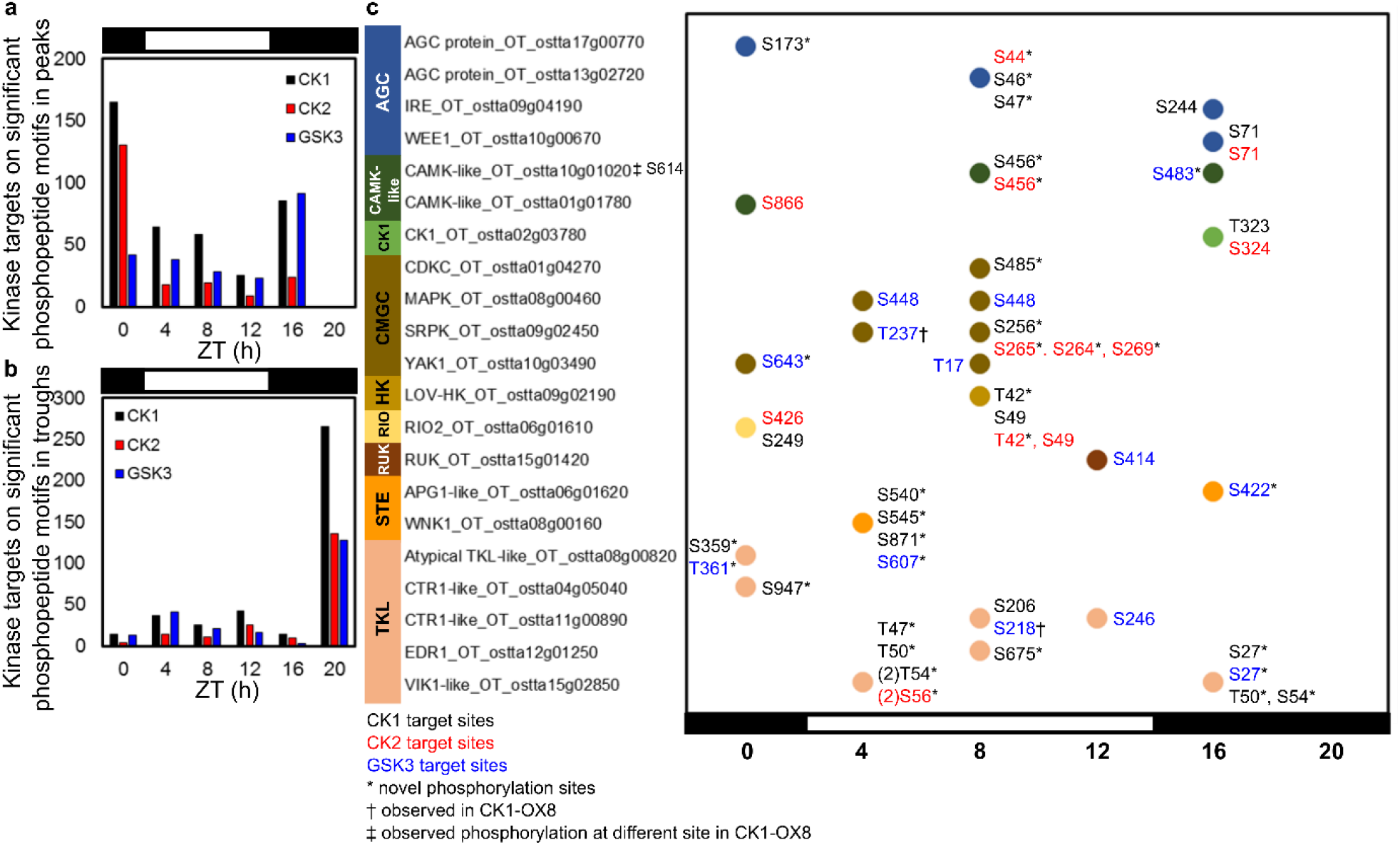
CK1, CK2 and GSK3 kinase targets and phosphorylation sites in rhythmic kinases. Distribution of GPS3-predicted CK1 (black), CK2 (red) and GSK3 (black) targets among rhythmic phosphopeptide motifs, binned by peak (a) and trough (b) times. (c) Phosphosites on rhythmic protein kinases predicted to be phosphorylated by CK1, CK2 and GSK3, site location labels coloured as in (a). * sites first reported here; †‡ sites observed previously (van Ooijen *et al*., 2013). Protein kinase classes are coloured as in Figure 5.

**Supplementary Figure S11.**
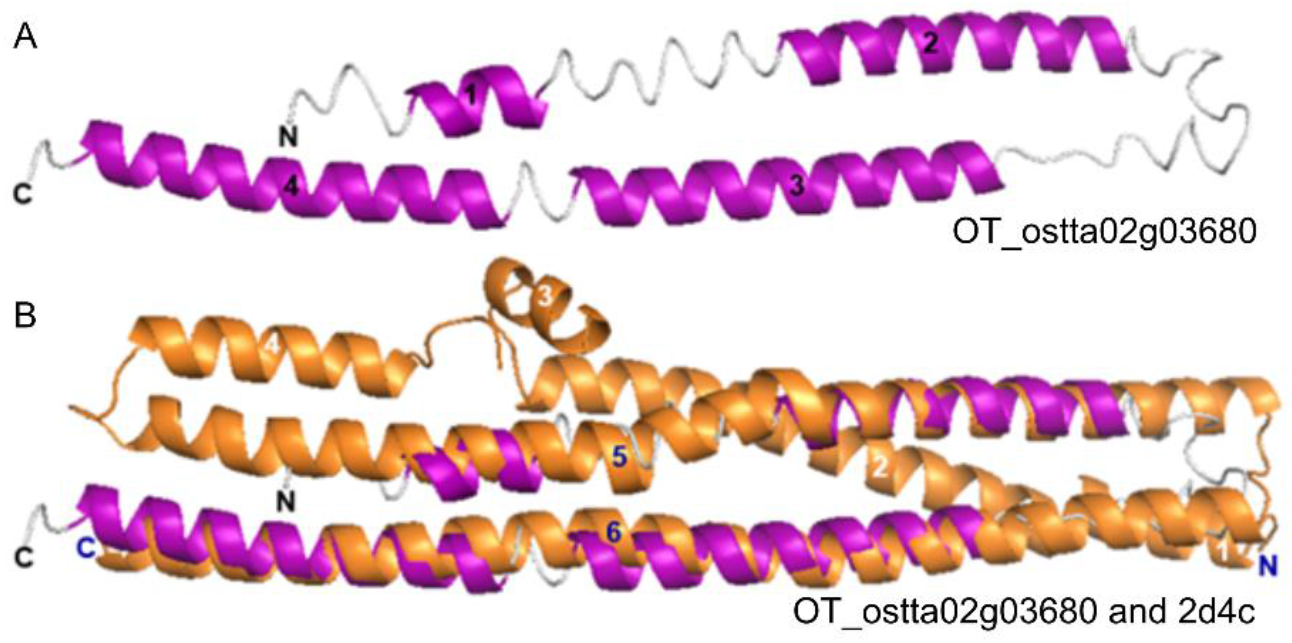
Structural homology of a rhythmic prasinophyte-specific protein. Structural homology models predicted using I-TASSER of (a) ostta02g03680 where the model is overlaid with (b) *H. sapiens* BAR domain structure (2d4c). Model α-helices (purple) and β-sheets (green) are numbered in black on the *O. tauri* model and in blue where structure is conserved with homologue protein overlay and in white where secondary structure is not conserved.

## Notes

http://dx.doi.org/10.6019/PXD001735

http://dx.doi.org/10.6019/PXD001734

http://dx.doi.org/10.6019/PXD002909

### Competing Interest Statement

The authors have declared no competing interest.

### Summary of Updates

We have clarified the main text, with a revised and updated Introduction, revised Results, and added Discussion section. In particular, overlapping proteomics results of Smallwood et al. bioRxiv 2018 are now discussed. Figure and Supplementary Figure panels are very similar or identical, with some rearrangements to fit the revised Results. The Methods section is little altered. Supplementary Tables are identical. Data Availability is updated.

